# A Newly Identified Class of Protein Misfolding in All-atom Folding Simulations Consistent with Limited Proteolysis Mass Spectrometry

**DOI:** 10.1101/2022.07.19.500586

**Authors:** Quyen V. Vu, Ian Sitarik, Yang Jiang, Divya Yadav, Piyoosh Sharma, Stephen D. Fried, Mai Suan Li, Edward P. O’Brien

## Abstract

Several mechanisms intrinsic to a protein’s primary structure are known to cause monomeric protein misfolding. Coarse-grained simulations, in which multiple atoms are represented by a single interaction site, have predicted a novel mechanism of misfolding exists involving off-pathway, non-covalent lasso entanglements, which are distinct from protein knots and slip knots. These misfolded states can be long-lived kinetic traps, and in some cases are structurally similar to the native state according to those simulations. Here, we examine whether such misfolded states occur in long-time-scale, physics-based all-atom simulations of protein folding. We find they do indeed form, estimate they can persist for weeks, and some have characteristics similar to the native state. Digestion patterns from Limited Proteolysis Mass Spectrometry are consistent with the presence of changes in entanglement in these proteins. These results indicate monomeric proteins can exhibit subpopulations of misfolded, self-entangled states that can explain long-timescale changes in protein structure and function *in vivo*.

**One-Sentence Summary:** Entangled misfolded states form in physics-based all-atom simulations of protein folding and have characteristics similar to the native state.

---

Several mechanisms intrinsic to a protein’s primary structure are known to cause monomeric protein misfolding (Table 1). Recently, high-throughput, coarse-grained simulations of protein synthesis and folding of the *E. coli* proteome have suggested there exists an additional widespread mechanism of misfolding^1–3^: proteins can populate off-pathway misfolded states that involve a change in non-covalent lasso entanglements^1,2^. This type of entanglement is defined by the presence of two structural components: a loop formed by a protein backbone segment closed by a non-covalent native contact, and another protein segment that is threaded through and around this loop^4–6^ in some cases multiple times (Figs. 1a and 1b). One-third of protein domains have such entanglements present in their native state^6^, and since most proteins are multidomain, around 70% of globular proteins contain these structural motifs.

**Table 1.**
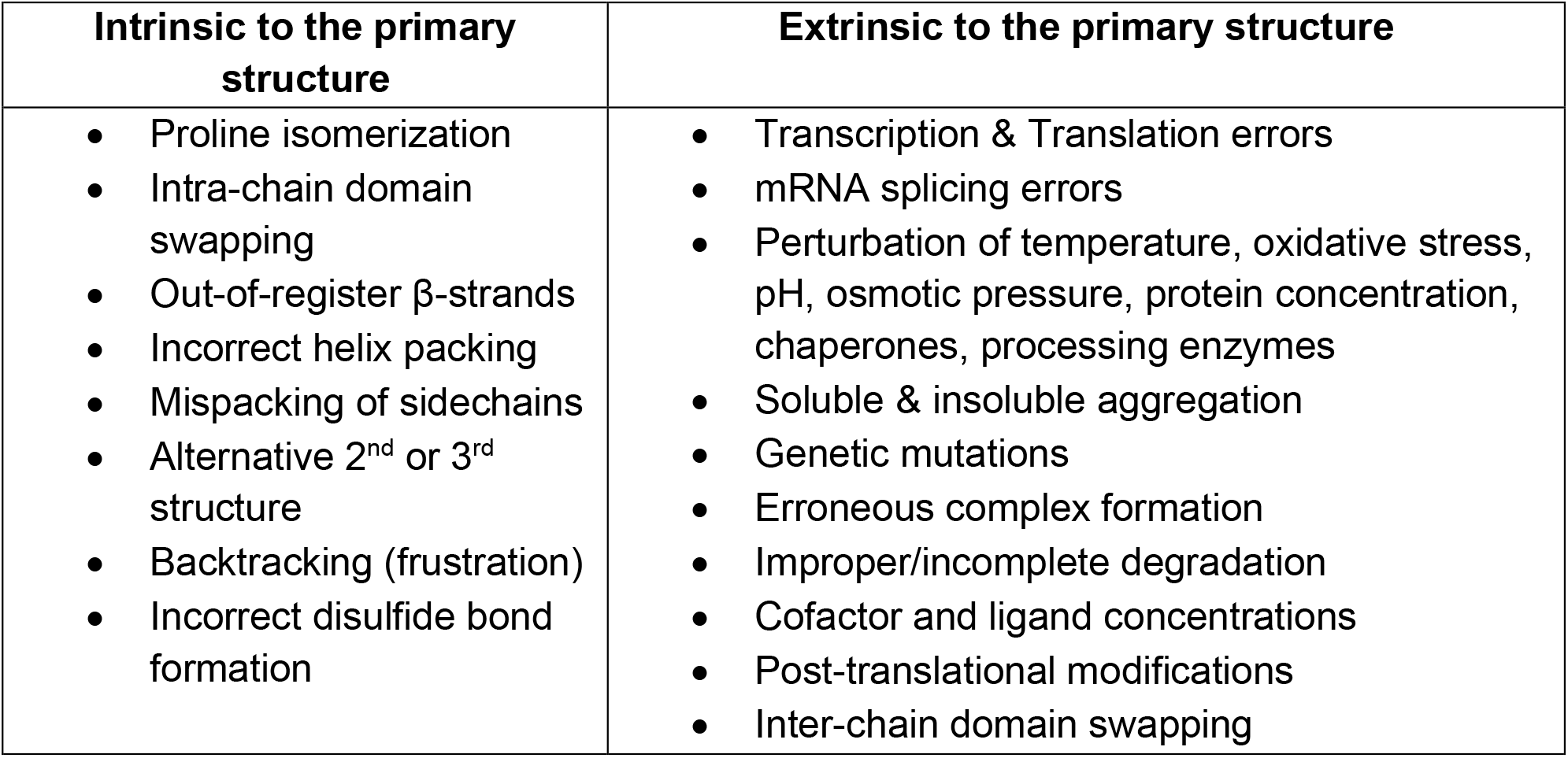
Mechanisms of protein misfolding^24–29^

| **Intrinsic to the primary structure** | **Extrinsic to the primary structure** |
| --- | --- |
| • Proline isomerization<br>• Intra-chain domain swapping<br>• Out-of-register β-strands<br>• Incorrect helix packing<br>• Mispacking of sidechains<br>• Alternative 2<sup>nd</sup> or 3<sup>rd</sup> structure<br>• Backtracking (frustration)<br>• Incorrect disulfide bond formation | • Transcription & Translation errors<br>• mRNA splicing errors<br>• Perturbation of temperature, oxidative stress, pH, osmotic pressure, protein concentration, chaperones, processing enzymes<br>• Soluble & insoluble aggregation<br>• Genetic mutations<br>• Erroneous complex formation<br>• Improper/incomplete degradation<br>• Cofactor and ligand concentrations<br>• Post-translational modifications<br>• Inter-chain domain swapping |

**Fig. 1.**
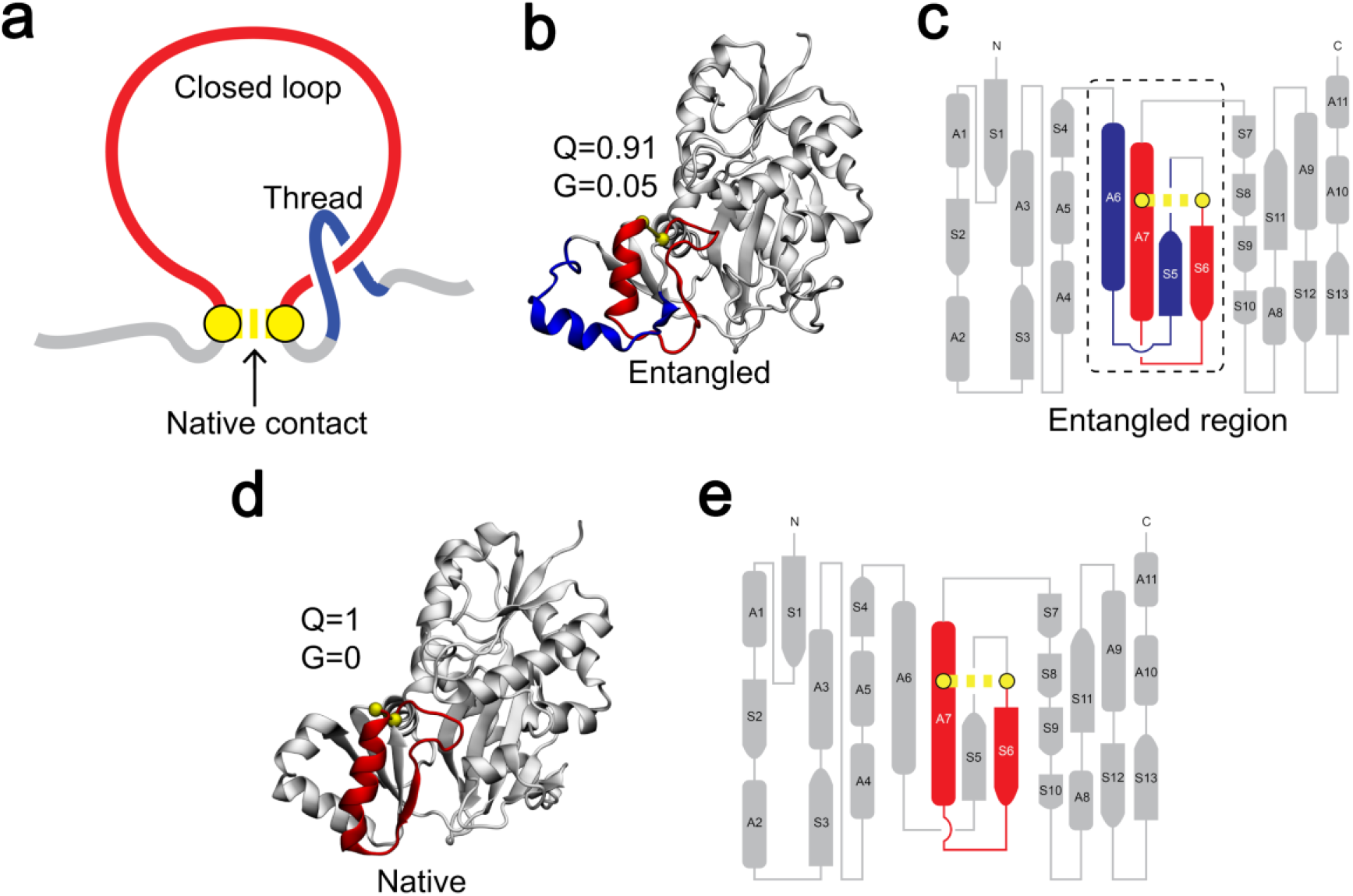
Visualizing misfolded changes of entanglement through various representations. (a) An illustration of the two geometric elements that compose an entanglement: the closed loop is colored in red and the threading segment is in blue. The loop is closed by a non-covalent contact between two residues (yellow). Such entanglements occur naturally in the native states of some proteins. In other cases, non-native formation of these entanglements can occur. (b) 3-D structure of a misfolded entangled state from the protein D-alanine—D-alanine Ligase B (DDLB; the closed loop and crossing section of the threading segment of their entangled regions are colored in red and blue, respectively) taken from previously reported coarse-grained simulations^1^. *Q* and *G* correspond, respectively, to the fraction of native contacts formed in the structure, and the fraction of native contacts that exhibit a change in entanglement. (c) A flattened 2-d structure representation of the entangled state of DDLB. S = beta sheet, A = alpha helix. (d) A 3-D structure of the crystal structure of DDLB. All native contacts are formed in the crystal structure (*Q*=1) and there is no entanglement change (hence, *G* = 0). (e) A flattened 2-d structure representation of the native state of DDLB. The coloring scheme of the relevant elements is the same in all panels.

The misfolded states observed in the aforementioned simulations involved either the gain of a non-native entanglement (*i*.*e*., the formation of an entanglement that is not present in the native ensemble, Table S1 and Fig. S1) or the loss of a native entanglement (*i*.*e*., an entanglement present in the native state fails to form, Table S1 and Fig. S1)^1–3^. This newly predicted class of misfolding offers an explanation for the decades-old observations that non-functional (or less functional) protein molecules can persist for long timescales without aggregating, and in the presence of chaperones neither get repaired nor degraded^7–9^.

The coarse-grained simulations, in which individual residues are represented by single interaction sites, suggest these misfolded states are often long-lived because to correctly fold they need to change their entanglement state: *e*.*g*., an entangled state would need to disentangle, which is energetically costly as already folded portions of the protein would need to unfold (see folded gray segments in Figs. 1b and 1c). Moreover, these states are likely to bypass cellular quality control mechanisms and remain long-lived in vivo^7–9^ because in some cases they are similar in size, shape, and structure to the native state (compare Figs. 1b and 1d).

Two criticisms of these findings are that they are based on a model with limited spatial resolution and use force field approximations that have the potential to affect the results. It could be the case, for example, that these states are never populated in a physics-based all-atom model. Further, the coarse-grained force field is “structure-based”, meaning the native state is encoded to be favored over other states^10^. Both of these approximations have the potential to affect these earlier findings.

Here, we examine whether a more fundamental, physics-based, all-atom model of the protein folding process exhibits these self-entangled states; and if they do, we test whether those states are off-pathway long-lived traps with properties similar to the native state. To do this, we analyze previously reported all-atom protein folding trajectories of ubiquitin^11^ and N-terminal domain (NTD) of λ-repressor^12^ starting from their native and unfolded states (Fig. 2a and b). Ubiquitin is a small protein (76 residues) found in eukaryotic organisms and regulates a range of processes including protein degradation and the cell cycle. Ubiquitin folds on the millisecond (ms) timescale under typical conditions *in vitro*^13^ and in about 3 ms in molecular dynamics simulations^11^. The NTD of λ-repressor in the published trajectories^12^ is 80 residues in length, binds DNA, and folds on the microsecond timescale^12,14^. Non-covalent lasso entanglements are not present in the native structure of either of these proteins.

**Fig. 2.**
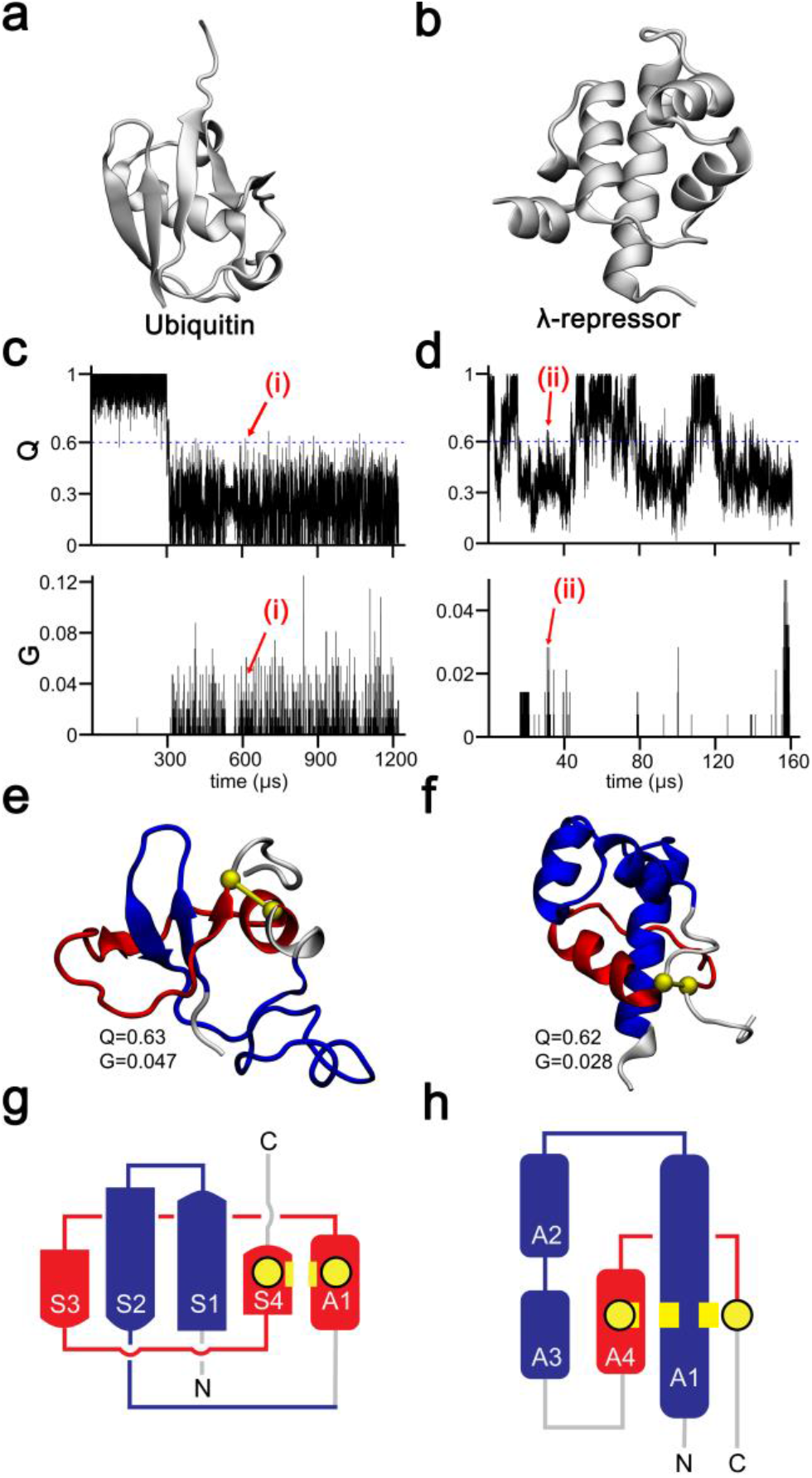
Misfolded gains-of-entanglement are observed in all-atom protein folding simulations. (a) Ubiquitin’s native structure and (b) λ-repressor’s native structure have no entanglement present. (c) Upper panel: Ubiquitin’s fraction of native contacts *Q* versus time from all-atom simulations, the blue-dashed line represents *Q* = 0.6. Lower panel: Ubiquitin’s degree of entanglement *G* versus time for the same trajectory. (d) Same as Fig. 2c, but for λ-repressor. (e) 3-D structure of a Ubiquitin entangled state observed at the time point labeled (i) in Fig. 2c, with entanglement elements colored as in Fig. 1a. (f) Same as Fig. 2e, but for λ-repressor, and from point labeled (ii) in Fig. 2d. (g) A flattened secondary structure representation for the entangled state in Fig. 2e. (h) Same as Fig. 2g, but for λ-repressor.

We analyzed whether non-native entanglements are populated during the folding trajectories of Ubiquitin and λ-repressor (Figs. 2c and d) totaling 8 milliseconds and 643 microseconds of simulation time, respectively. We observed several non-native entangled states (denoted by high, non-zero *G* values (eq. S4) in Figs. 2c and d) during folding transitions (indicated by shifts from low to high *Q* values in Figs. 2c and d). Thus, protein self-entangled states are observed in all-atom models. Next we asked whether these entangled states are off-pathway. We never observe in these trajectories a direct transition from a (high *G*, low *Q*) pair of values to a (low *G*, high *Q*) pair of values from one time point to the next; meaning there is never a direct transition from the misfolded state to the native state. Hence, these misfolded states are off pathway – they must unfold to reach the native state. These entangled states are clearly short-lived (lasting just a few nanoseconds according to Figs. 2c and d), seemingly at odds with the prediction that these can be kinetically trapped states^2^. However, these simulations were performed at high temperatures (390 K for Ubiquitin and 350 K for λ-repressor), near their melting temperatures in the all-atom force field and configurational transitions are accelerated. Therefore, these states may be able to rapidly disentangle due to these high temperatures.

To test if these states are kinetic traps at physiological temperatures we performed unrestrained molecular dynamics simulations at 310 K (37 °C). We selected entangled structures that had at least 60% of their native contacts (*Q*) formed as starting structures, resulting in 21 and 12 structures, respectively, for Ubiquitin and λ-repressor. Structurally, these entangled states are qualitatively similar (Table S2), 20 out of the 21 ubiquitin structures have closed-loops located towards the C-terminus and the threading segment composed of the N-terminus. For λ-repressor, the loop forms towards the C-terminal end, and is very similar in all conformations, and the N-terminal threads through it. Representative structures for these entangled states are shown in Figs. 2e and f. Three independent molecular dynamics trajectories were started from each of these conformations and the simulations were run for 700 ns, or until the entanglement was lost (*i*.*e*., *G* = 0 was reached).

For the 63 trajectories (= 21 × 3) of ubiquitin, 71% (45 out of 63) contain a non-native entanglement that persists for 700 ns (Fig. S2a and Table S2). On average then, the time to disentangle is approximately 2.1 μs (estimated using eq. S6, setting *t* = 700 ns). For λ-repressor, 17% of trajectories (6 out of 36) persist in an entangled state for 700 ns (Fig. S2b and Table S2). The average time to disentangle for λ-repressor is approximately 390 ns (estimated using eq. S6 setting *t* = 700 ns), 50 times faster than for ubiquitin entanglements.

Relative to their time-scale of protein folding, these entangled states are not long-lived kinetic traps even at 310 K. Based on the previous coarse-grained simulation results this is to be expected^1^, since ubiquitin and NTD are small single-domains proteins. They are representative of the “model proteins” that have been traditionally studied by biophysicists and which demonstrate the ability to rapidly and efficiently unfold and refold^15^, but are not representative of the complexity of proteomes^16^, (e.g., the median protein length in yeast is 379 residues^17^). In such very small proteins, entanglements compose a large proportion of the total protein structure present – and hence, it is easier to disentangle as most of the protein structure is misfolded and less stable. For example, for the entanglements reported in Table S2, 79% of ubiquitin’s primary structure is involved in the entanglement (*i*.*e*., either part of the loop or threading segment, Fig. 2g), and 83% of λ-repressor’s primary structure is involved in the entanglement (Fig. 2h). In contrast, a protein of median length with a misfolded entanglement will need to unfold substantial amounts of already folded structure to permit disentanglement (Figs. 3a-f). Unfolding for these proteins is energetically more costly, and hence near-native entanglements in large proteins are more likely to be long-lived kinetic traps^2^.

**Fig. 3.**
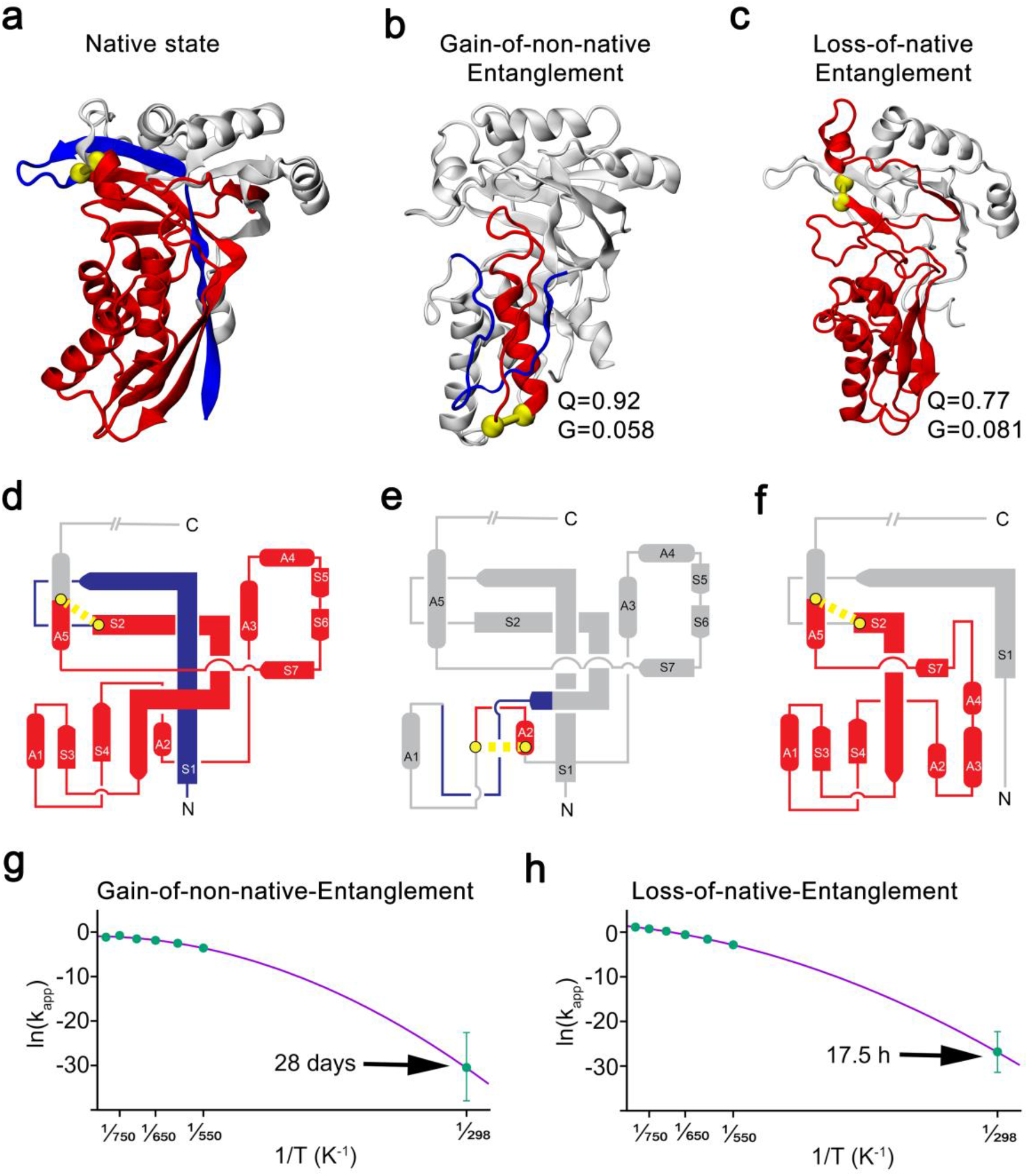
Misfolded states involving either the gain of a non-native entanglement or loss of a native entanglement in a larger protein are estimated to persist from hours to weeks. (a) Native structure of 4-Diphosphocytidyl-2-C-Methyl-D-Erythritol Kinase (PDB ID: 2WW4). (b) Gain of a non-native entanglement structure in this protein. 92% of native contacts are present in this structure (Q=0.92). (c) Loss of a native entanglement structure. 77% of native contacts are present in this structure (Q=0.77) and 8% of the native contacts exhibit a change in entanglement state relative to the native state (G=0.081). (d)-(f) Flattened secondary structure representation of the structures shown in panels (a)-(c). Structure in C-terminal is shown as a simple line as it does not involve the native entanglement. (g) Arrhenius plot for the gain of a non-native entanglement state, with a super-Arrhenius fit^22,23^, error bars of the natural logarithm of disentanglement rate at high temperatures are smaller than symbols, and the extrapolated lifetime of disentanglement at 298 K is about 28 days (corrected for the acceleration in unfolding rates observed in all-atom simulations). Error bars represent the 95% confidence interval about the mean. K_app_ in a unit of ns^-1^. (h) Same as Fig. 3g but for the loss of native entanglement state, the time it takes to unfold is extrapolated to between 2 and 18 hours (Table S4), the plot shows the results for *Q*_*threshold*_ = 0.2.

Unrestrained all-atom simulations are incapable of folding such large proteins on tractable simulation time scales. Therefore, to test this “size-effect” hypothesis we calculated the lifetime of two misfolded states of a larger protein, *E. coli*’s 4-Diphosphocytidyl-2-C-Methyl-D-Erythritol Kinase (Fig. 3a; 283 residues; gene ispE). This protein was chosen from our dataset of proteins that exhibit entanglements in coarse-grained simulations^10^ because of its large size and because it exhibits both classes of misfolding: a conformation with a gain of a non-native entanglement and another with a loss of a native entanglement (Figs. 3b,c and Table S3). We carried out *in silico* temperature jump experiments on both of these misfolded states of this enzyme that were created by back-mapping coarse-grained conformations to an all-atom representation. Arrhenius analyses were then carried out (Figs. 3g and 3h), corrected for the observed acceleration in unfolding rates in all-atom simulations (see SI Methods Section 4), to estimate the lifetime of this entangled state at 298 K (this temperature was used to allow comparison to experimental unfolding data measured at this temperature; see SI Method for details). We calculate that it takes 28 days for the gain of a non-native entanglement misfolded state to disentangle, a necessary step to reach the folded state (Fig. 3g and Table S4, specifically, 2.413×10^6^ s (95% Confidence Interval: [931.587 s, 4.338×10^9^ s])). For the misfolded state involving the loss of a native entanglement, we calculate the time it takes for this off-pathway state to unfold, a necessary step to reach the folded state, is between 2 and 18 hours (Fig. 3h, see Table S4 for statistics). Thus, misfolded changes of entanglement can be very long-lived states in larger proteins according to physics-based all-atom simulations.

To experimentally test for the existence of these misfolded states, we unfolded purified 4-Diphosphocytidyl-2-C-Methyl-D-Erythritol Kinase (ispE) in 6 M guanidinium chloride, initiated refolding by dilution, and then probed the protein’s conformation with Limited Proteolysis Mass Spectrometry^18,19^ (LiP-MS, see SI Methods). LiP-MS allows us to pinpoint subtle structural differences between folded and misfolded conformations^2,20^. Based on the predicted timescales from simulations, we allowed ispE three relatively long refolding times (1 h, 10 h, and 24 h, Table S8 and Supplementary Data) before probing it with limited proteolysis. Specifically, we examined if any of the predicted changes in structure accompanying self-entanglement overlap with the proteolysis products that exhibit a significant change in abundance in the refolded samples relative to native ispE. If there was significant overlap at 24 hrs, and at least one other time point, we considered these predicted changes in self-entanglement to be consistent with the mass spec data (Table S8, bolded).

In the LiP-MS data we observe two positions with perturbed proteolytic susceptibility after both 10 h and 24 h of refolding: residue 210 (an aspartic acid) and residue 161 (a threonine). From this, we conclude that structural differences persist for at least 24 hours for some structural subpopulations of this kinase. Next we cross referenced these two cut sites against our entangled states (Table S8), and found they overlap with the structural changes observed in the aforementioned ‘loss of entanglement’ conformation (Fig. 3c). Specifically, crossing residue 16, in the native entanglement, is in van der Waals contact with residue 210 in the native state. When this native entanglement fails to form in the misfolded state, it increases the solvent exposure of residue 210 by 5%, making residue 210 more susceptible to protease digestion (Fig. 4a). Consistently, residue 210 experiences a modest (3-4–fold) but significant (p<0.01 by t-test with Welch’s correction, Supplementary Data) increase in proteolytic susceptibility. Crossing residue 33, which forms a non-native entanglement in this misfolded state, forms a van der Waals contact with residue 165 – a neighbor to the cut site 161 – and increases residue 161’s solvent exposure by 28% (Fig. 4b). The probability of this agreement between these mass spec data and simulated entanglements arising by random chance is 1 in 53 (p-value = 0.019, see Methods). Thus, this misfolded state is highly consistent with this LiP-MS digestion pattern. The ‘gain of non-native entanglement’ misfolded state (Fig. 3b) exhibits a significant overlap with the LiP-MS data only at the 1 hour time point (cut site residue 118 (Table S8) is in contact with the crossing residue 44, Fig. 4c). Therefore, there is structural agreement, but this misfolded state may persist in reality for less than 10 hours rather than the 28 days estimated by the simulations. Overall, we see generally consistent agreement between the predicted structural changes accompanying entanglement and experimental data (Table S8 and Supplementary Data).

**Fig. 4:**
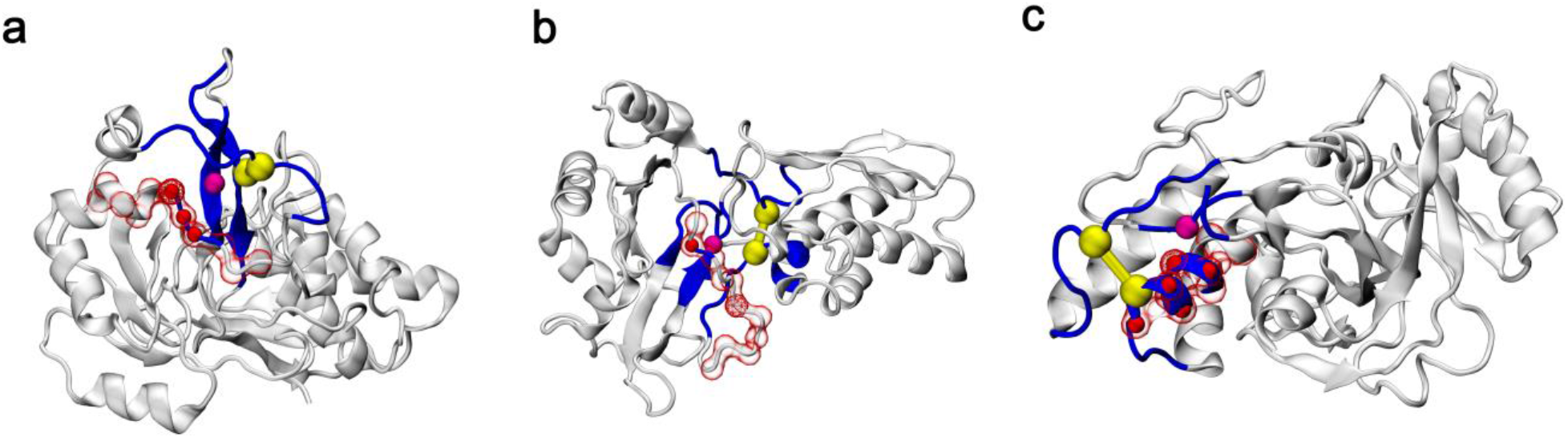
Overlap between the LiP-MS Proteinase K cut sites and the observed misfolded structures in 4-Diphosphocytidyl-2-C-Methyl-D-Erythritol Kinase. (a-c) The two residues that form the native contact to close the loop, the residue that crosses the plane of the loop on the threading segment, and the backbone segment making the loop are shown, respectively, as two yellow spheres, a magenta sphere, and a blue segment. The remaining portions of the protein structure are shown in gray. The residue that is cut by Proteinase K is shown as a red-mesh sphere, along with +/-5 of its nearest neighboring residues along the primary structure are shown in red-transparent volume. The residues overlapping between these entangled residues and the Proteinase K cut site are shown as red-solid spheres. (a) The loss-of-native-entanglement structure (same conformation as in Fig. 3c, shown in different viewpoint) is consistent with the Proteinase K cut site at residue D210, observed in the mass spec data at the 10 and 24-hour time points (Table S8). (b) Same conformation as in Fig. 3c (loss-of-native-entanglement structure, shown from a different viewpoint) but has contact gains of non-native entanglement with entangled residues overlap with PK cut site at residue T161, observed in the mass spec data at the 10 and 24-hour time points (Table S8). (c) A gain-of-non-native entanglement structure (same conformation as in Fig. 3b, shown from a different viewpoint) illustrating the overlap with Proteinase K cut site L118, observed in the mass spec data at the 1-hour time point (Table S8). Note well, the bar connecting the two yellow spheres is emphasizing there is a non-covalent interaction; there is not a disulfide bond between these two residues.

Finally, we analyzed how native-like are these three protein’s entangled states. The radius of gyration of these states are comparable to the native state (the differences are less than 10%, Table S5), indicating the entangled and native states are of similar sizes. These entangled states are estimated to have a solubility similar to the native state (Table S5) according to a model that accounts for the chemical properties of the exposed protein surface^1^. This indicates these misfolded states are likely to remain as soluble in solution as the native fold. Next, we analyzed the secondary structure of proteins using Stride software^21^ and find that the non-native entangled states contain up to 80% of the secondary structure found in the native state (Table S2). Thus, the size, solubility, and secondary structure analyses indicate these misfolded states have similarity to the native state. This is again consistent with LiP-MS data for ispE in that only 2–4 peptides (out of >100) displayed significant differences in susceptibility at the different refolding times. In other words, ispE can spontaneously refold much of its native structure, with changes in entanglement resulting in subtle, local changes.

In conclusion, non-native self-entanglements occur during protein folding in all-atom models, some of these states are long-lived, and have properties similar to the native state. The simulated misfolded states are consistent with those populated during refolding from denaturant and probed by protease susceptibility. This new class of protein misfolding opens up novel avenues of research including new targets for drug design, expanding our understanding of protein homeostasis in cells, and the impact of these states on protein function.

## Acknowledgments

We thank D.E Shaw’s group for providing access to their MD trajectories.

## Data and materials availability

Full details of all-atom molecular dynamics simulations, analyses, and Limited Proteolysis Mass Spectrometry methods are included in the supporting information (PDF). Summary data for these experiments are also provided in Supplementary Data (XLSX). Code used to analyze entanglement is available at https://github.com/obrien-lab-psu/entanglement_analysis_public. Software used in experimental data analysis is available at https://github.com/FriedLabJHU/Refoldability-Tools.

## References

1. Jiang, Y. et al. How synonymous mutations alter enzyme structure and function over long time scales. Nat. Chem. (2022) In press.

2. Nissley, D. A. et al. Universal protein misfolding intermediates can bypass the proteostasis network and remain soluble and less functional. Nat. Commun. 13, 3081 (2022).

3. Halder, R., Nissley, D. A., Sitarik, I. & O’Brien, E. P. Subpopulations of soluble, misfolded proteins commonly bypass chaperones: How it happens at the molecular level. bioRxiv 2021.08.18.456736 (2021) doi:10.1101/2021.08.18.456736.

4. Baiesi, M., Orlandini, E., Trovato, A. & Seno, F. Linking in domain-swapped protein dimers. Sci. Rep. 6, 1–11 (2016).

5. Baiesi, M., Orlandini, E., Seno, F. & Trovato, A. Exploring the correlation between the folding rates of proteins and the entanglement of their native states. J. Phys. A Math. Theor. (2017).

6. Baiesi, M., Orlandini, E., Seno, F. & Trovato, A. Sequence and structural patterns detected in entangled proteins reveal the importance of co-translational folding. Sci. Rep. 9, 1–12 (2019).

7. Komar, A. A., Lesnik, T. & Reiss, C. Synonymous codon substitutions affect ribosome traffic and protein folding during in vitro translation. FEBS Lett. 462, 387–391 (1999).

8. Zhou, M. et al. Non-optimal codon usage affects expression, structure and function of clock protein FRQ. Nature 494, 111–115 (2013).

9. Zhou, M., Wang, T., Fu, J., Xiao, G. & Liu, Y. Nonoptimal codon usage influences protein structure in intrinsically disordered regions. Mol. Microbiol. 97, 974–987 (2015).

10. Nissley, D. A. et al. Electrostatic Interactions Govern Extreme Nascent Protein Ejection Times from Ribosomes and Can Delay Ribosome Recycling. J. Am. Chem. Soc. 142, 6103–6110 (2020).

11. Piana, S., Lindorff-Larsen, K. & Shaw, D. E. Atomic-level description of ubiquitin folding. Proc. Natl. Acad. Sci. U. S. A. 110, 5915–5920 (2013).

12. Lindorff-Larsen, K., Piana, S., Dror, R. O. & Shaw, D. E. How fast-folding proteins fold. Science (80-.). 334, 517–520 (2011).

13. Sivaraman, T., Arrington, C. B. & Robertson, A. D. Kinetics of unfolding and folding from amide hydrogen exchange in native ubiquitin. Nat. Struct. Biol. 8, 331–333 (2001).

14. Yang, W. Y. & Gruebele, M. Folding at the speed limit. Nature 423, 193–197 (2003).

15. Maxwell, K. L. et al. Protein folding: Defining a “standard” set of experimental conditions and a preliminary kinetic data set of two-state proteins. Protein Sci. 14, 602–616 (2005).

16. Braselmann, E., Chaney, J. L. & Clark, P. L. Folding the proteome. Trends Biochem. Sci. 38, 337–344 (2013).

17. Brocchieri, L. & Karlin, S. Protein length in eukaryotic and prokaryotic proteomes. Nucleic Acids Res. 33, 3390–3400 (2005).

18. Feng, Y. et al. Global analysis of protein structural changes in complex proteomes. Nat. Biotechnol. 32, 1036–1044 (2014).

19. To, P., Whitehead, B., Tarbox, H. E. & Fried, S. D. Nonrefoldability is Pervasive across the E. coli Proteome. J. Am. Chem. Soc. 143, 11435–11448 (2021).

20. To, P., Xia, Y., Lee, S. O. Devlin, T., Fleming, K. G. & Fried, S. D. A proteome-wide map of chaperone-assisted protein refolding in a cytosol-like milieu. Proc. Natl. Acad. Sci. USA. (2022) In press.

21. Frishman, D. & Argos, P. Knowledge-based protein secondary structure assignment. Proteins Struct. Funct. Bioinforma. 23, 566–579 (1995).

22. Onuchic, J. N., Luthey-Schulten, Z. & Wolynes, P. G. THEORY OF PROTEIN FOLDING: The Energy Landscape Perspective. Annu. Rev. Phys. Chem. 48, 545–600 (1997).

23. Sarkar, D., Kang, P., Nielsen, S. O. & Qin, Z. Non-Arrhenius Reaction-Diffusion Kinetics for Protein Inactivation over a Large Temperature Range #. ACS Nano 13, 8669–8679 (2019).

24. Gosavi, S., Chavez, L. L., Jennings, P. A. & Onuchic, J. N. Topological frustration and the folding of interleukin-1β. J. Mol. Biol. 357, 986–996 (2006).

25. Sherman, M. Y. & Qian, S. B. Less is more: Improving proteostasis by translation slow down. Trends Biochem. Sci. 38, 585–591 (2013).

26. Kiefhaber, T., Kohler, H. H. & Schmid, F. X. Kinetic coupling between protein folding and prolyl isomerization. I. Theoretical models. J. Mol. Biol. 224, 217–229 (1992).

27. Kikis, E. A. The intrinsic and extrinsic factors that contribute to proteostasis decline and pathological protein misfolding. Advances in Protein Chemistry and Structural Biology vol. 118 (Elsevier Ltd, 2019).

28. Salahuddin, P., Siddiqi, M. K., Khan, S., Abdelhameed, A. S. & Khan, R. H. Mechanisms of protein misfolding: Novel therapeutic approaches to protein-misfolding diseases. J. Mol. Struct. 1123, 311–326 (2016).

29. Lafita, A., Tian, P., Best, R. B. & Bateman, A. Tandem domain swapping: determinants of multidomain protein misfolding. Curr. Opin. Struct. Biol. 58, 97–104 (2019).

