## Supplementary Information for "A Newly Identified Class of Protein Misfolding in All-atom Folding Simulations Consistent with Limited Proteolysis Mass Spectrometry"

### Supplementary Method

#### 1. All-atom simulations.

Initial conformations of the protein were placed at the center of a rectangular periodic box with the minimum distance between the box edge and protein atoms of 1.5 nm, the system was solvated with TIP3P water model[1], and NaCl was added to neutralize and mimic relevant salt concentrations (Ubiquitin: 0 mM, Lambda-repressor: 50 mM, *E. Coli* 4-Diphosphocytidyl-2c-Methyl-D-Erythritol Kinase: 150 mM, the salt concentration was added according to original studies[2,3]). Energy minimization was then carried out using the steepest descent algorithm[4,5] to minimize steric conflict. The systems were then gradually heated from 2 K to 310 K over 1 ns. After that, the system was equilibrated for 1 ns in NVT and 1 ns NPT ensemble with all heavy atoms restrained using a harmonic potential with a force constant of 1000 kJ/mol/nm$^2$ before it went into production run in the NPT ensemble. The Particle Mesh Ewald method[6] was used to calculate long-range electrostatic interactions beyond 12 Å and the Lennard-Jones interactions were calculated with a cut-off distance of 12 Å and applied smoothly switching the forces to zero between 10 Å and 12 Å. Temperature and pressure were maintained at 310 K and 1 atm using Nose-Hoover thermostat[7,8] and Parrinello-Rahman barostat[9], respectively. The LINCS algorithm[10] was used to constrain all bonds involving hydrogen atoms and the integration time step was set to 2 fs. Simulations were performed using GROMACS 2018[11] with the CHARMM36m forcefield[12]. This procedure was used in all the new simulations carried out in this study.

#### 2. Characterizing changes in entanglement.

Calculating changes in entanglement has been reported elsewhere[13–15]. Here, we briefly describe our procedure. To detect non-covalent lasso entanglements we use linking numbers, which to compute requires at least one closed loop. We define this loop as being composed of the backbone trace connecting residues $i$ and $j$ that form a non-covalent contact in the given protein conformation. The looped portion (red segment in Fig. 1a) is identified if the $C_\alpha$ coordinates of residues $i$ and $j$ (yellow spheres in Fig. 1a) are closer than 9 Å and $|i-j| \geq 10$ residues. Outside this loop is an N-terminal segment, composed of residues 1 through $i-1$, and a C-terminal segment composed of residues $j+1$ through $N$. These two segments represent open curves, whose entanglement through the closed loop was characterized by linking numbers denoted $g_N$ and $g_C$. For a given structure of an $N$ residues protein, with a contact present at residues $(i, j)$, the average coordinates $\boldsymbol{R}_l$ and the gradient d$\boldsymbol{R}_l$ of the point $l$ on the curves were calculated as:

$$\begin{cases} \boldsymbol{R}_l = \frac{1}{2}(\boldsymbol{r}_l + \boldsymbol{r}_{l+1}) \\ d\boldsymbol{R}_l = \boldsymbol{r}_{l+1} - \boldsymbol{r}_l \end{cases} \tag{S1}$$

where $\boldsymbol{r}_l$ is the coordinates of the $C_\alpha$ atom in residue $l$. The linking numbers $g_N(i, j)$ and $g_C(i, j)$ of N- and C-terminal, respectively, were calculated as:

$$\begin{cases} \boldsymbol{g}_N(i,j) = \dfrac{1}{4\pi}\displaystyle\sum_{m=6}^{i-5}\sum_{n=i}^{j-1}\dfrac{\boldsymbol{R}_m - \boldsymbol{R}_n}{|\boldsymbol{R}_m - \boldsymbol{R}_n|^3}(d\boldsymbol{R}_m \times d\boldsymbol{R}_n) \\ \boldsymbol{g}_C(i,j) = \dfrac{1}{4\pi}\displaystyle\sum_{m=i}^{j-1}\sum_{n=j+4}^{N-6}\dfrac{\boldsymbol{R}_m - \boldsymbol{R}_n}{|\boldsymbol{R}_m - \boldsymbol{R}_n|^3}(d\boldsymbol{R}_m \times d\boldsymbol{R}_n) \end{cases} \quad \text{(S2)}$$

where we excluded the first 5 residues on the N-terminal curve, the last 5 residues on the C-terminal curve, and 4 residues before and after the native contact to eliminate the error introduced by both the high flexibility and contiguity of the termini and trivial entanglements in local structure. It is worth noting that partial linking values between 0.5 – 0.7 may not signify real entanglements and thus those have to be checked manually[16]. The above integrations yield two non-integer values, and the total linking number for a contact ($i$, $j$) is estimated as the sum of N- and C-terminal linking numbers:

$$g(i,j) = round[g_N(i,j)] + round[g_C(i,j)] \quad \text{(S3)}$$

Comparing the absolute value of the total linking number for a contact ($i$,$j$) at a given conformation to that of a reference state (*i.e.*, native state) allows us to ascertain a gain or loss of linking between the backbone trace loop and the terminal open curves as well as any switches in chirality. There are 6 changes in linking cases we should consider (Table S1 and Fig. S1) when using this approach to quantify entanglement.

We defined the G metric, degree of entanglement, as a time-dependent order parameter that reflects the extent of the topological entanglement changes in a given structure compared to the native structure and is calculated as:

$$G(t) = \frac{1}{M}\sum_{(i,j)}\Theta\left[(i,j) \in NC \bigcap g(i,j,t) \neq g^{native}(i,j)\right] \quad \text{(S4)}$$

where ($i$,$j$) is one of the native contacts in the native crystal structure; *NC* is the set of native contacts formed in the current structure at time $t$; $g(i,j,t)$ and $g^{\text{native}}(i,j)$ are, respectively, the total linking number of the contact ($i$,$j$) at time $t$, and native structures estimated using Eq. S3; *M* is the total number of native contacts within the native structure and the selection function $\Theta$ equals 1 when the condition is true and equals 0 when it is false. The larger $G$ is, the greater the number of native contacts that have changed their entanglement status relative to the reference state.

##### 3. Disentanglement time constant from unrestrained simulations at room temperature.

The disentangled time constant can be estimated through the connection with the portion of disentangled trajectories:

$$P_{dis}(t) = 1 - e^{\frac{-t}{\tau}} \tag{S5}$$

$$\tau = \frac{-t}{ln[1 - P_{dis}(t)]} \tag{S6}$$

Where, $P_{dis}(t)$ is the portion of disentangled trajectories at time $t$. $\tau$ is a time constant that entanglement disappears. We use this relation to estimate the average lifetime of entangled states of Ubiquitin and λ -repressor by setting t = 700 ns.

##### 4. In silico Temperature jump simulations, and Arrhenius analysis of 4-Diphosphocytidyl-2-C-Methyl-D-Erythritol Kinase

For the 'gain of non-native entanglement' misfolded state, we calculate the time it takes for the entangled misfolded state to disentangle, permitting proper folding. The temperatures studied were 550K, 600K, 650K, 700K, 750K, and 800K. We perform 30 trajectories in the NVT ensemble for each temperature, then get the list of first passage times that there is no gain in entanglement in structure compared to the reference state, which is: $G_0+G_1 = 0$ ($G_0$ and $G_1$ are two kinds of gain in entanglement defined in Table S1). To take into account the fluctuation in $G$ parameters, we consider there is no gain in entanglement in structure if $G_0+G_1 = 0$ for 1 ns. The survival probability was then fitted to a single-exponential function to find the fitting parameters ($t_0$, k):

$$S_U(t) = \begin{cases} 1, & 0 \leq t < t_0 \\ e^{-k(t-t_0)}, & t \geq t_0 \end{cases} \tag{S7}$$

Where $t_0$ is the delay time of entanglement (Fig. S4). The apparent rate at temperature $T$ is: $k_{app}(T) = \frac{1}{t_0+\frac{1}{k}}$

The apparent rates are found to have a super-Arrhenius behavior[17,18] and are then fitted as the function of $T$: $\ln(k_{app}) = \frac{a}{T^2} + \frac{b}{T} + c$, and we then extrapolate to the target temperature 298K, where a, b, c are the free fitting parameters.

The disentanglement time in simulation is: $\tau_{sim} = \frac{1}{k_{app}}$.

To map the disentanglement time from simulation time to real-time we scale the disentanglement time by the factor $\alpha = \frac{k_{unfolding}(simulation)}{k_{unfolding}(experiment)}$, which accounts for observed differences between experimental versus simulated unfolding rates. Ideally, $\alpha$ would be computed for each protein individually, under the same environmental conditions in the experiment, and using the same force field. However, due to the lack of experimental measurement of ispE's unfolding rate and extra computational cost, we utilize the scaling factor from the unfolding process of the protein DHFR, from another study[13]. In that study, the authors used the AMBER ff14SB force field (while we use CHARMM36m) and the same Arrhenius procedure described here and compared to its experimental unfolding rate at 298 K to find the scaling factor $\alpha = 143$ . That is, the unfolding rate in the all-atom simulations are 143 times faster than in reality. Thus, we multiplied the disentangling characteristic time $\tau_{sim}$ by 143.

For the 'loss of a native entanglement' misfolded state, we calculate the time it takes for this off-pathway state to unfold using the same procedure but for each temperature, we performed 50 statistically independent trajectories. Unfolding criteria is $Q \leq Q_{threshold}$ for 1 ns with various values of $Q_{threshold}$ to determine the unfolding time. The results are presented in Table S4, and Fig. S5 shows for $Q_{threshold} = 0.2$.

#### 5. Secondary structure similarity definition.

The secondary structure similarity in Table S2 is defined as the fraction of residues that are in the correct secondary element at the current structure compared to the native structure, we only count residues in alpha-helix or beta-sheet. The secondary structural elements of protein are assigned by the STRIDE program[19].

#### 6. Prediction of the solubility of the entangled structures.

Solubility of the entangled structures in Table S5 is defined as the percent soluble protein estimated via the insolubility propensities of structure $i$ ($\chi_i^{sol}$), fully disordered structure, which has no secondary and tertiary structure elements ($\chi_{disordered}^{insol}$), and the minimum propensity values of all structures ($\min(\chi_i^{sol})$):

$$f_i^{sol} = \frac{\chi_{disordered}^{insol} - \chi_i^{insol}}{\chi_{disordered}^{insol} - \min(\chi_i^{insol})} \quad \text{(S8)}$$

The insolubility was estimated by taking into account the aggregation propensity ($\chi^{agg}$), degradation propensity ($\chi^{deg}$), and minus the Hsp70 binding propensity ($\chi^{Hsp70}$) that is considered to prevent the misfolding protein from aggregation:

$$\chi^{insol} = \chi^{agg} + \chi^{deg} - \chi^{Hsp70} \quad \text{(S9)}$$

For a given entangled structure, the aggregation, degradation, and Hsp70 binding propensities were estimated as the relative change of the solvent-accessible surface area (SASA) of the aggregation-prone, degradation-prone, and Hsp70 binding regions of the entangled state $i$ against the SASA of the native structure:

$$\chi_i^T = \frac{SASA_i^T}{SASA_{native}^T} \quad \text{(S10)}$$

Where $T$ can be aggregation, degradation, or Hsp70 binding propensities. The aggregation-prone region was predicted by the AmylPred2 server[20]. The degradation-prone region was defined as the hydrophobic residues (Ile, Val, Phe, Cys, Met, Ala, Gly, and Trp). The Hsp70 binding region was predicted by using ChaperISM[21].

#### 7. Preparation of ispE Native and Refolded Samples for LiP-MS

Recombinant 4-diphosphocytidyl-2-C-methyl-D-erythritol (CDP-MEP) kinase (ispE) from *Salmonella typhimurium* with a C-terminal His-tag, purified from *E. coli* was procured from Echelon Biosciences, USA as frozen concentrated stocks (8.89 mg/mL). The native samples were prepared by diluting1 µL of protein stock 350-fold with a 49:1 mixture of native buffer (20 mM Tris pH 8.2, 100 mM NaCl, 2 mM $MgCl_2$, 1 mM DTT) and unfolding buffer (20 mM Tris pH 8.2, 100 mM NaCl, 2 mM $MgCl_2$, 7 M GdmCl, 10 mM DTT). The native samples were prepared in triplicate. To unfold the samples, 2.5 µL of ispE protein stock was diluted 7-fold with unfolding buffer and incubated overnight at room temperature. The next day, unfolded samples were refolded by dilution 50-fold with native buffer. The samples were refolded in triplicates at three-time points (1 h, 10 h, and 24 h).

A proteinase K (PK, from *Tritirachium album*) stock solution of 0.032 µg/µL was prepared in a 1:1 mixture of native buffer and 20% glycerol). The stock of PK was stored at -20 ºC and thawed at most once before use. Limited proteolysis was performed by adding 250 µL of native and refolded proteins samples (1 h, 10 h, and 24 h) to 2 µL of PK stock (enzyme:substrate 1:100 *w/w*) and incubated at room temperature for 1 min. The PK activity was quenched by placing the sample tubes in a mineral oil bath equilibrated at 105 ºC for 5 min.

#### 8. MS Sample Preparation

Boiled proteolyzed samples were transferred to a fresh microfuge tube containing 216 mg of urea and 35 µL of a native buffer such that the final volume of solution was 450 µL at 8 M urea concentration. Samples were reduced by the addition of dithiothreitol (DTT) to 10 mM final concentration and incubation at 37 ºC with agitation (700 rpm) in a benchtop ThermoMixer (Eppendorf) for 45 minutes. Samples were then alkylated by adding iodoacetamide (IAA) to 40 mM final concentration and incubated at room temperature in the dark for 30 min. Following alkylation, samples were diluted 4-fold with 50 mM ammonium bicarbonate pH 8 (final concentration of urea 2 M) and digested with trypsin (1:50 enzyme:protein *w/w* ratio, Pierce) overnight (ca. 16 h) at 25 ºC, 700 RPM in a ThermoMixer. Protein digests were desalted using 1 cc Sep-Pak C18 cartridges (Waters) over a vacuum manifold.

Digested peptides were acidified by adding trifluoroacetic acid (TFA, Acros) to a final concentration of 1% (*v/v*). Sep-Pak cartridges were briefly conditioned (2 × 1 mL of 80%

acetonitrile, 0.5% TFA in Optima water) and equilibrated (4 × 1 mL 0.5% TFA in Optima water). Acidified digests were loaded onto the cartridges under a reduced vacuum (such that it took approximately 5 min for 1 mL to passage through the sorbent) and then washed (4 × 1 mL 0.5% TFA in Optima water). The cartridges were placed on top of 15 mL conical tubes, and 1 mL of Buffer B was pipetted onto the resin. The cartridges were gently eluted under centrifugal force in swing-buckets spun at 350 rpm in a 5910R centrifuge (Eppendorf) for 3 min and then reduced to dryness in a Vacufuge centrifugal concentrator (Eppendorf) and stored at -80 ºC until further use.

#### 9. LC-MS/MS acquisition

This study utilized an UltiMate3000 UHPLC system coupled with a Q-Exactive HF-X Orbitrap mass spectrometer (Thermo Fisher). Peptide fractions were resuspended in 30 µL of 0.1% Optima formic acid and 2% acetonitrile in Optima water, vigorously vortexed, and sonicated for five minutes. The amount of peptide material in each resuspended sample was approximately quantified with a NanoDrop One$^{C}$ microvolume UV-Vis spectrophotometer (Thermo Fisher Scientific), and typically ~0.5 µg of the peptide was injected for each run. The column and trap cartridge temperatures were set to 40 ºC, and the flow rate was set to 300 µL/min. Mobile phase Solvent A comprised 0.1% FA, 2% acetonitrile, and 98% Optima water. Mobile phase Solvent B comprised 0.1% FA in 98% acetonitrile, and 2% Optima water.

The injected sample was loaded onto a trap column cartridge (Acclaim PepMap 100, C18, 75 µm × 2 cm, 3 µm, 100 Å column) and washed with Solvent A for 10 minutes. Next, the trap column was switched to be in line with the separating column (Acclaim Pepmap RSLC, C18, 75 µm × 25 cm, 2 µm, 100 Å column). The sample was separated in a linear gradient from 2% B to 35% B over 100 minutes, 40% B over 25 minutes, and 90% B over 5 minutes. The end of the LC method employed a saw-tooth gradient to remove sample residue from the resolving column.

A full MS scan was acquired in positive ion mode at a mass range of 350–1500 *m/z*. The resolution of the full MS scans was set to 120k, the automatic gain control (AGC) target was set to 8E3, and the maximum injection time was set to 64 ms. After each full MS scan, twenty data-dependent MS$^2$ scans were collected at a resolution of 15k, an AGC target of 1E5, a minimum AGC target of 8E3, a maximum injection time of 55 ms, and an isolation window of 1.4 *m/z.* To dissociate precursors prior to their reanalysis by MS$^2$, peptides were subjected to an HCD of 28% normalized collision energies. Fragments with charges of 1, 6, 7, or higher and unassigned were excluded from analysis, and a dynamic exclusion window of 30.0 s was used for the data-dependent scans.

#### 10. LC-MS/MS data analysis

Proteome Discoverer (PD) Software Suite (v2.4, Thermo Fisher) and the Minora Algorithm were used to analyze mass spectra and perform Label-Free Quantification (LFQ) of detected peptides[14,22,23]. Default settings for all analysis nodes were used except where specified. The data were searched against ispE from *Salmonella typhimurium* (P30753, Uniprot[24]). The PD Spectrum Files RC node was used for peptide identification, using a semi-tryptic search allowing up to 2 missed cleavages. A precursor mass tolerance of 10 ppm was used for the MS1 level, and a fragment ion tolerance was set to 0.02 Da at the $MS^2$ level. Peptide lengths between 6 and 144 amino acid residues were allowed with a peptide mass between 350 and 5000 Da. Oxidation of methionine and acetylation of the N-terminus were allowed as dynamic modifications, while carbamidomethylation on cysteines was set as a static modification. The Fixed Value PSM validator node was used for FDR validation. Raw normalized extracted ion intensity data for the identified peptides were exported from the .pdResult file using a three-level hierarchy (protein > peptide group > consensus feature). These data were further processed utilizing custom Python analyzer scripts (available on GitHub and described in depth previously[25,26]). Briefly, normalized ion counts were collected across the refolded replicates, and the native replicates for each successfully identified peptide group. Effect sizes are the ratio of averages (reported in $\log_2$), and *P*-values (reported as $-\log_{10}$) were assessed using t-tests with Welch's correction for unequal population variances. Missing data are treated in a special manner. If a feature is not detected in all three native (or refolded) injections and is detected in all three refolded (or native) injections, we use those data and fill the missing values with 1000 (the ion limit of detection for this mass analyzer); this peptide becomes classified as an all-or-nothing peptide. The missing value is dropped if a feature is not detected in one of six injections. Any other permutation of missing data (e.g., missing in two injections) results in the quantification getting discarded. We consider only those peptides with a greater than 2-fold difference in abundance in the refolded sample versus the native sample ($log_2(R/N) \geq 1$, column E in Supplementary Data) and whose difference is statistically significant ($p \leq 0.01, -log_{10}(P) \geq 2$, column F in Supplementary Data).

#### 11. Determination of representative changes in entanglements

We k-means clustered the last 100 ns of coarse-grained post-translational simulations resulting from 50 independent synthesis simulations[14,27] along two order parameters that capture the nativeness of the structures (fraction of native contacts, Q) and the changes in self-entanglement of the protein (fraction of native contacts with a change in self-entanglement, G). These microstates are then coarse-grained into metastable states based on the PCCA++ algorithm[28] (Figure S3). A structure was chosen at random from the 5 most probable microstates in each metastable state and analyzed for changes in entanglement as described in Supplementary Method Section 2. We clustered the raw changes in entanglement together based on the number of times a thread pierces the loop plane, and the rounded partial linking numbers and change types of each termini (Table S1). Within each of these clusters, we selected the entanglement with the minimal loop as a representative entanglement. Since we had a finite set of structures representing the most probable microstates in each metastable state we checked the clustering by hand to ensure cluster internal homogeneity and uniqueness.

#### 12. Overlap of representative change in entanglement with LiP-MS data

Given a set of representative entanglements $E_R$, a set of significant half-tryptic LiP-MS peptides $L_{s(t)}$ at various time points $t \in \{1hr, 10hr, 24hr\}$ we calculate the extent of overlap, $M_i$, for each representative entanglement $E_{R(i)} \in E_R$ as

$$M_i = \sum_t \frac{1}{|L_{s(t)}|} \sum_k J(E_{R(i)}, L_{s(t,k)}) \quad \text{(S11)}$$

Where $J(E_{R(i)}, L_{s(t,k)})$ is the jaccard similarity between the set of entangled residues $E_{R(i)}$ for a representative entanglement $i$ and the set of residues within ±5 residues of the Proteinase K cut site observed for a given LiP-MS peptide $L_{s(t,k)} \in L_{s(t)}$

$$J(E_{R(i)}, L_{s(t,k)}) = \frac{|E_{R(i)} \cap L_{s(t,k)}|}{|E_{R(i)} \cup L_{s(t,k)}|} \quad \text{(S12)}$$

where $k$ is an arbitrary index for the peptide. We define the entangled residues of a representative entanglement as the residues within 5 amino acids of the native contacts forming the loop and residues within 8 Å of the residues that cross the loop plane determined by Topoly[29] (*i.e.* thread the loop). We then define a rank-ordered vector for the entanglements that have non-zero overlap at the longest time point and at least one other time point as $M \equiv [M_{1,i}, M_{2,i}, \dots, M_{n,i}]$ were $M_{1,i}, \geq M_{2,i} \geq \cdots \geq M_{n,i}$. We finally only consider the top two entanglements $\alpha$ and $\beta$ that have the greatest overlap as $M(\alpha, \beta) = [M_{1,\alpha}, M_{2,\beta}]$. We can define a similar metric for comparison of each representative entanglement $E_{R(i)} \in E_R$ with a set of randomly drawn peptides $L'_{s(t,r)}$ from the set of all the possible peptides identified in the LiP-MS refolding experiment for the protein $L$ and denote the top overlaps as $M'_r(\alpha, \beta)$. Where $r$ is a dummy variable denoting the index of a random sample if multiple random samples are being tested. The *p-value* is then calculated by:

$$p = \frac{1}{|R|} \sum_{r=1}^{|R|} \Upsilon(r) \quad \text{(S13)}$$

Where,

$$\Upsilon(r) = \begin{cases} 0 & if \quad M'_r(\alpha, \beta) < M(\alpha, \beta) \\ 1 & if \quad M'_r(\alpha, \beta) \geq M(\alpha, \beta) \end{cases} \quad \text{(S14)}$$

and $R$ is a super-set of 100000 sets of permutated peptides chosen from all potentially possible peptides $L'_{s(t,r)}$ and $\Upsilon(r)$ is an indicator function that determines whether the top two observed significant overlaps are less than the top two test significant overlaps for one of the permutated sets in $R$.

### Supplementary Figures

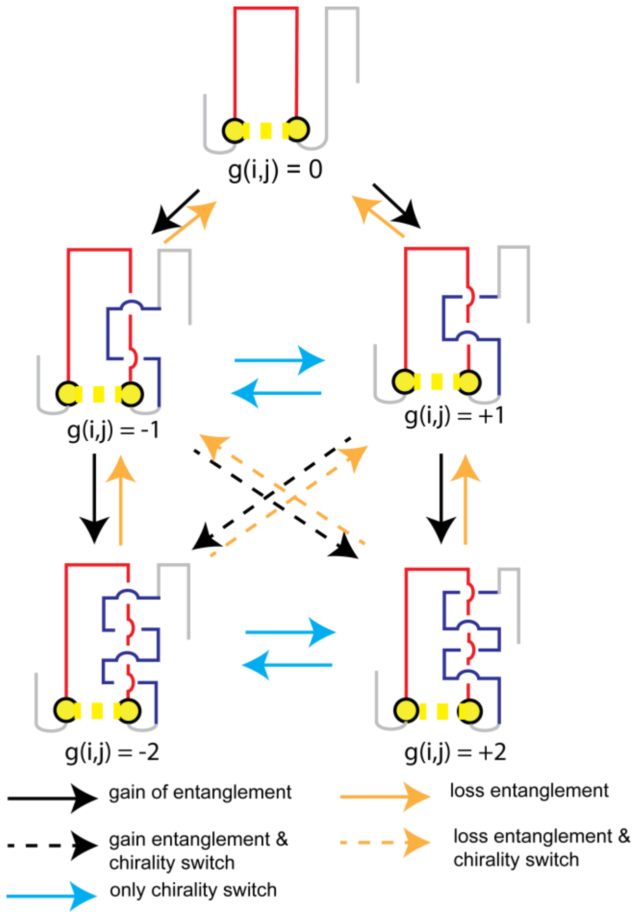


**Figure S1: Schematic of different changes in topological entanglement observed with the Gaussian linker integration method.** The closed loop is colored in red and the threading segment is in blue. The loop is closed by a non-covalent contact between two residues (yellow). The total linking number $g(i,j)$ of contact present at residues $(i, j)$ provides information on the topological entanglement between a loop formed by a non-covalent contact and the flanking terminal thread.

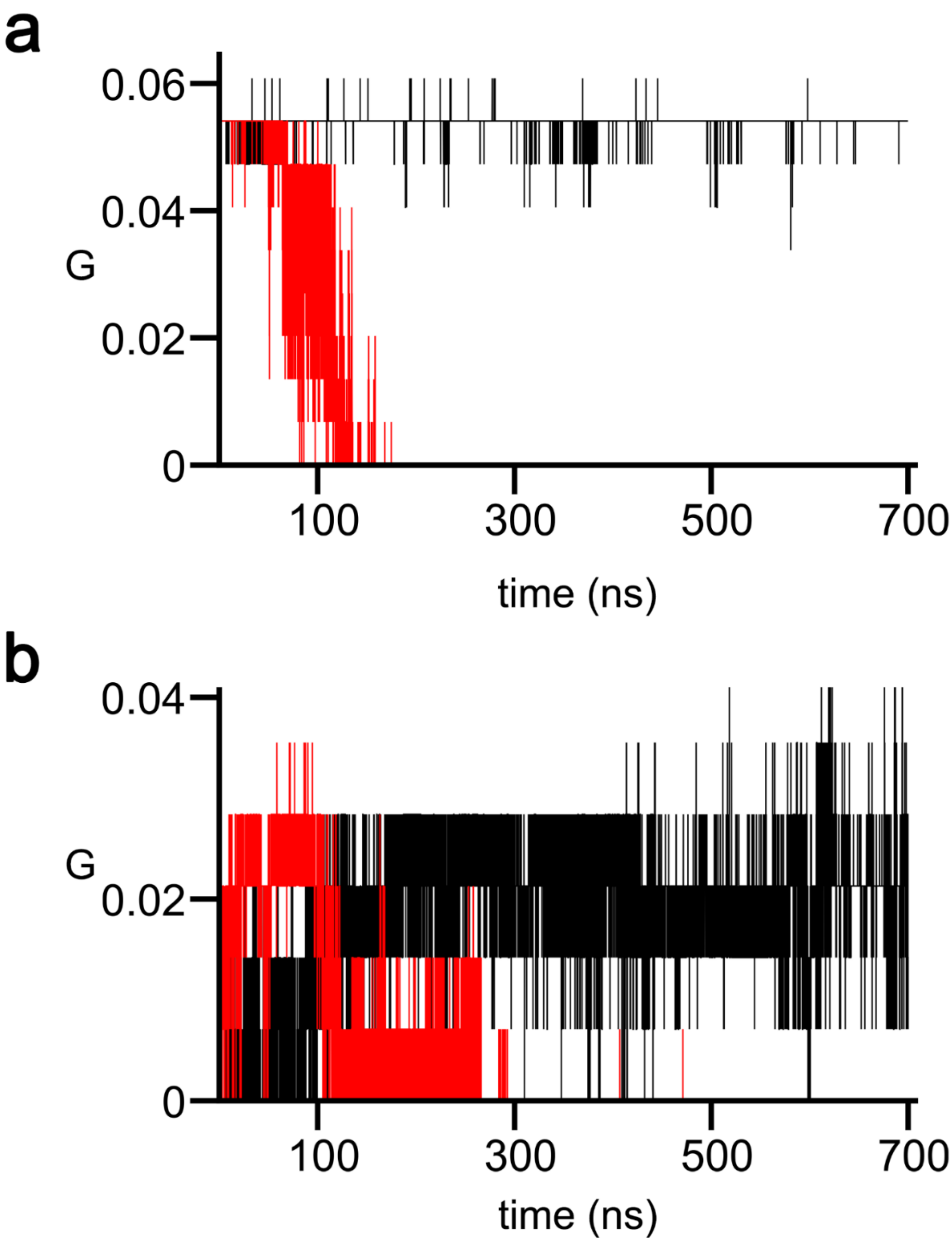


**Figure S2:** (a) Ubiquitin's G versus time for two trajectories, one trajectory does not disentangle up to 700 ns (black line) and the other trajectory disentangles at a time less than 700 ns (red line). (b) Same as Fig. S2a, but for λ-repressor.

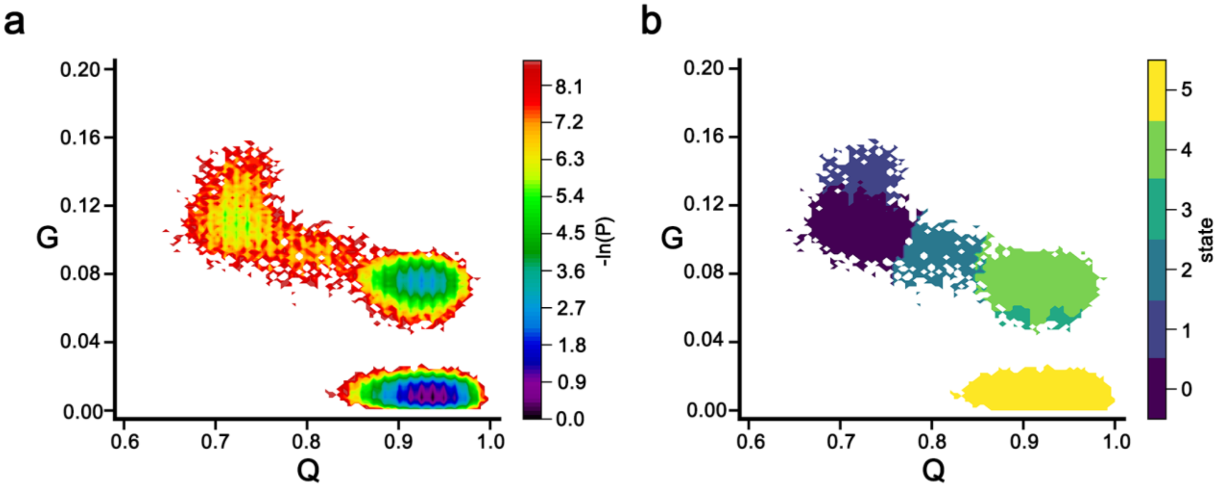


**Figure S3:** (a) -ln(P) surface of the last 100ns of post-translational time of the coarse-grained synthesis simulations of 4-Diphosphocytidyl-2-C-Methyl-D-Erythritol Kinase demonstrating several metastable basins. (b) The metastable states after k-means clustering.

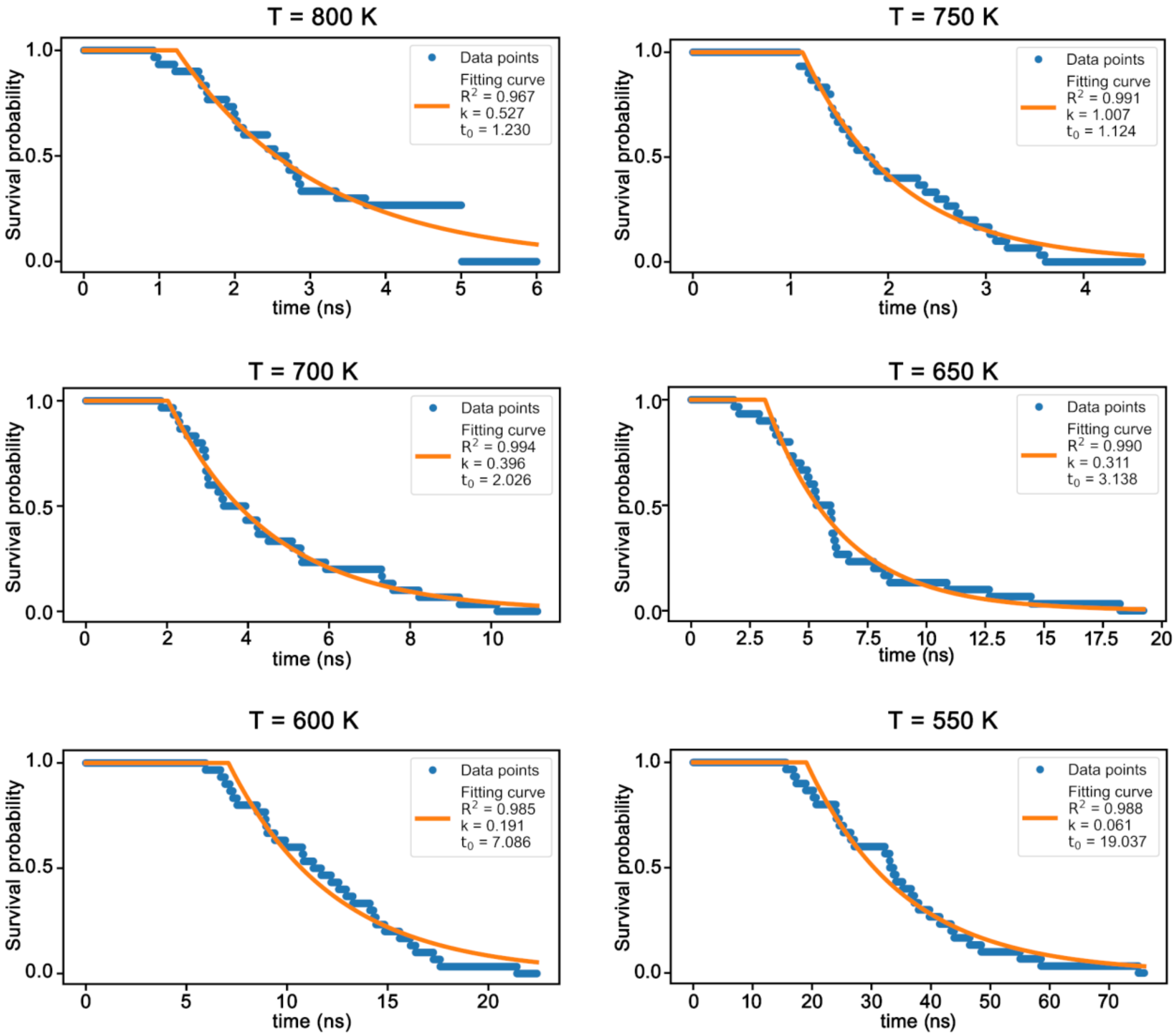


**Figure S4: Estimation of the disentanglement timescale for kinase misfolded gain of a non-native entanglement state.** Survival probabilities of entangled structures vs. time at different simulation temperatures that were fitted by an exponential function (orange). The coefficient of determination $R^2$ and fitted parameters (rate k and lag time $t_0$) are presented in the legends.

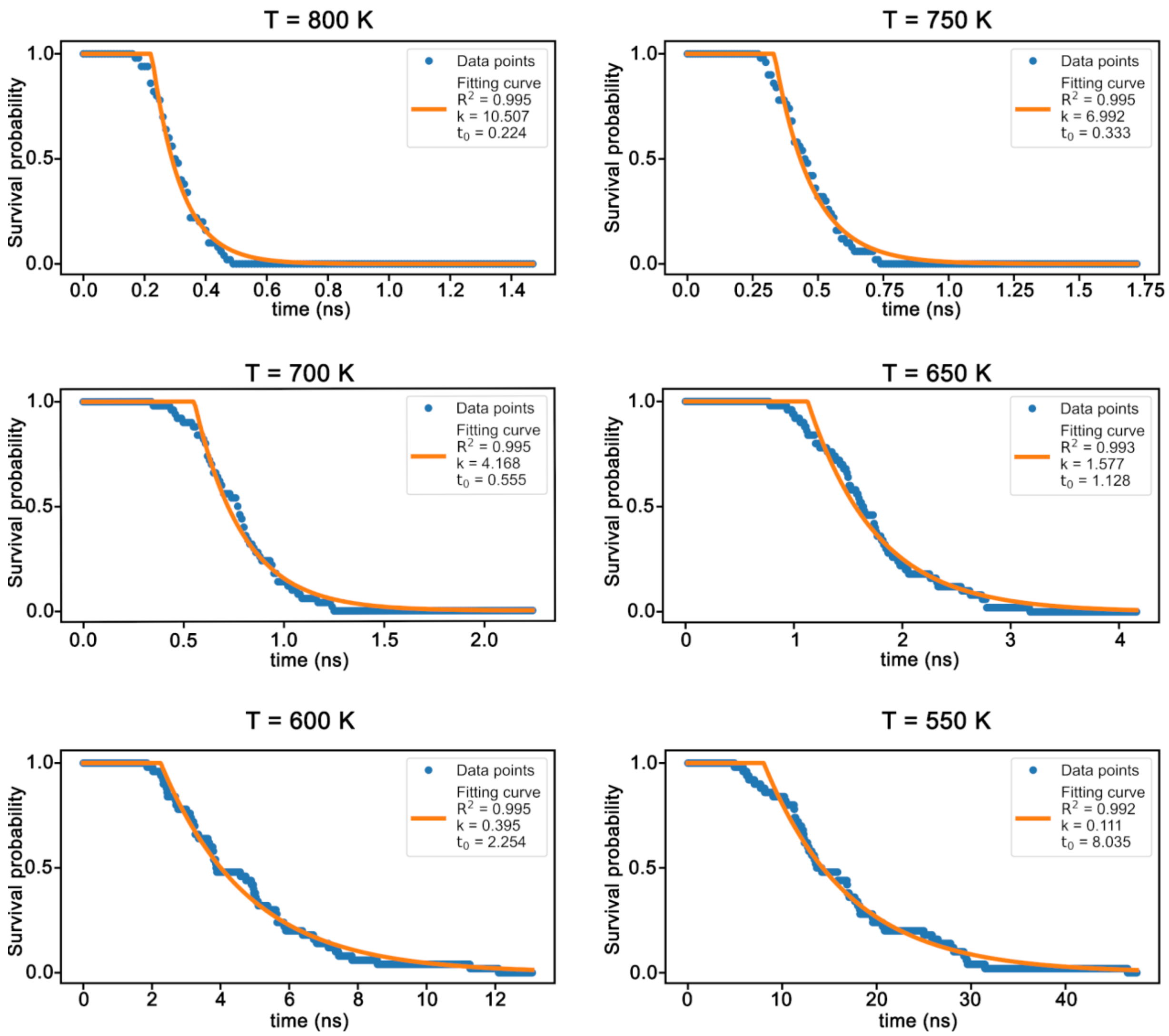


**Figure S5: Estimation of the unfolding timescale for kinase misfolded loss of native entanglement state.** An example plot for the threshold to define unfolded state $Q \leq 0.2$ for 1 ns. Survival probabilities of entangled structures vs. time at different simulation temperatures that were fitted by an exponential function (orange). The coefficient of determination $R^2$ and fitted parameters (rate k and lag time $t_0$) are presented in the legends.

### Supplementary Tables

**Table S1**: Definitions of the different types of change in entanglement that are possible.

| Type | Change in entanglement | Change in chirality | Conditions |
|---|---|---|---|
| $G_0$ | Gain | No | $\|g^{current}(i,j)\| > \|g^{native}(i,j)\|$ $and$ $g^{current}(i,j) \times g^{native}(i,j) \geq 0$ |
| $G_1$ | Gain | Yes | $\|g^{current}(i,j)\| > \|g^{native}(i,j)\|$ $and$ $g^{current}(i,j) \times g^{native}(i,j) < 0$ |
| $G_2$ | Lose | No | $\|g^{current}(i,j)\| < \|g^{native}(i,j)\|$ $and$ $g^{current}(i,j) \times g^{native}(i,j) \geq 0$ |
| $G_3$ | Lose | Yes | $\|g^{current}(i,j)\| < \|g^{native}(i,j)\|$ $and$ $g^{current}(i,j) \times g^{native}(i,j) < 0$ |
| $G_4$ | None | Yes | $\|g^{current}(i,j)\| = \|g^{native}(i,j)\|$ $and$ $g^{current}(i,j) \times g^{native}(i,j) < 0$ |
| $G_5$ | None | None | $\|g^{current}(i,j)\| = \|g^{native}(i,j)\|$ $and$ $g^{current}(i,j) \times g^{native}(i,j) \geq 0$ |

**Table S2:** Properties of entangled states of Ubiquitin and λ-repressor. The conformational ID was indexed by sorting the degree of entanglement G in descending order for each protein.

| ID (1) | Fraction of native contacts Q (2) | Degree of entanglement G (3) | Residues that close the loop and crossing residues (4) | Secondary structure similarity (5) | Persistence Time (ns) (6) |
|---|---|---|---|---|---|
| Ubiquitin | | | | | |
| 1 | 0.61 | 0.054 | [45-67], [5,13] | 0.71 | 700, 700, 520 |
| 2 | 0.64 | 0.054 | [45-67], [5,13] | 0.60 | 215, 700, 553 |
| 3 | 0.63 | 0.054 | [45-68], [5,13] | 0.57 | 700, 700, 481 |
| 4 | 0.61 | 0.054 | [44-69], [5,13] | 0.43 | 700, 700, 700 |
| 5 | 0.61 | 0.054 | [44-70], [5,12] | 0.33 | 700, 337, 700 |
| 6 | 0.63 | 0.054 | [45-67], [5,13] | 0.69 | 700, 163, 150 |
| 7 | 0.61 | 0.054 | [44-67], [5,13] | 0.64 | 700, 700, 627 |
| 8 | 0.61 | 0.054 | [44-70], [5,12] | 0.48 | 700, 700, 700 |

| | | | | | |
|---|---|---|---|---|---|
| 9 | 0.63 | 0.054 | [45-67], [5,13] | 0.52 | 700, 700, 578 |
| 10 | 0.61 | 0.054 | [45-67], [5,13] | 0.55 | 700, 700, 700 |
| 11 | 0.63 | 0.054 | [45-67], [5,13] | 0.57 | 700, 700, 700 |
| 12 | 0.63 | 0.054 | [45-67], [5,13] | 0.60 | 700, 464, 666 |
| 13 | 0.61 | 0.054 | [45-68], [5,12] | 0.50 | 700, 121,700 |
| 14 | 0.63 | 0.054 | [45-67], [5,13] | 0.48 | 700, 700, 700 |
| 15 | 0.63 | 0.054 | [45-68], [5,13] | 0.55 | 700, 700, 700 |
| 16 | 0.61 | 0.054 | [44-70], [5,13] | 0.60 | 700, 700, 700 |
| 17 | 0.63 | 0.054 | [44-70], [5,13] | 0.48 | 700, 700, 700 |
| 18 | 0.61 | 0.054 | [45-67], [5,12] | 0.50 | 700, 700, 700 |
| 19 | 0.63 | 0.054 | [45-68], [5,13] | 0.57 | 700, 700, 700 |
| 20 | 0.63 | 0.047 | [45-67], [4,13] | 0.19 | 40, 100, 55 |
| 21 | 0.74 | 0.034 | [23-53], [66] | 0.81 | 0, 0, 0 |
| λ-repressor | | | | | |
| 22 | 0.62 | 0.028 | [57-75], [9] | 0.71 | 1, 22, 1 |
| 23 | 0.61 | 0.028 | [57-75], [11] | 0.71 | 12, 18, 615 |
| 24 | 0.62 | 0.028 | [57-75], [11] | 0.73 | 1, 2, 25 |
| 25 | 0.61 | 0.021 | [57-75], [12] | 0.64 | 2, 700, 523 |
| 26 | 0.65 | 0.021 | [57-75], [11] | 0.75 | 700, 0, 182 |
| 27 | 0.62 | 0.021 | [54-75], [12] | 0.73 | 1, 19, 2 |
| 28 | 0.61 | 0.021 | [53-75], [11] | 0.73 | 178, 700, 200 |
| 29 | 0.61 | 0.014 | [57-75], [12] | 0.68 | 2, 3, 18 |
| 30 | 0.63 | 0.014 | [57-75], [11] | 0.73 | 106, 67, 603 |
| 31 | 0.65 | 0.007 | [57-75], [12] | 0.73 | 0, 217, 0 |
| 32 | 0.63 | 0.007 | [57-75], [12] | 0.71 | 700, 178, 0 |
| 33 | 0.66 | 0.007 | [54-75], [11] | 0.71 | 700, 700, 10 |

(1) Entangled state ID used in this study.

(2) Fraction of native contacts, we only selected entangled structures that had at least 60% of their native contacts formed as the starting structures for the subsequent all-atom simulations.

(3) Degree of entanglement G, is the fraction of native contacts with changes in entanglement calculated by Eq. S4.

(4) Representative native contact that closes the loop (first square brackets) and list of crossing residues (second square brackets). Note well, some entanglements have a threading segment that pierces the loop once (hence a single crossing residue is reported), while in others the threading segment pierces the loop twice (hence two crossing residues are reported).

(5) Secondary structure similarity is defined as the fraction of residues that are in their native secondary structure in the current structure.

(6) Time it takes for the non-native entanglement to disentangle. 3 replicas were run for each entangled state found from D.E. Shaw's protein folding trajectories. Note well that the simulations were only run for 700 ns. Therefore, the entries of 700 ns mean the entanglement stated formed the entire simulation time.

**Table S3.** Entanglement descriptions of 4-Diphosphocytidyl-2-C-Methyl-D-Erythritol Kinase starting structures in Arrhenius analysis. ($G_0$, $G_1$, $G_2$, $G_3$, and $G_4$) are the type of change in entanglement defined in Table S1, #NC is the total number of native contacts in the crystal structure, and equals 739. We refer to the structure by its dominant kind of change in entanglement.

| Structure | Q | $G = \frac{\sum_{i=0}^{4} G_i}{\# NC}$ | $G_0$ | $G_1$ | $G_2$ | $G_3$ | $G_4$ |
|---|---|---|---|---|---|---|---|
| Loss of native entanglement | 0.77 | 0.081 | 4 | 0 | 55 | 1 | 0 |
| Gain of non-native entanglement | 0.92 | 0.058 | 40 | 0 | 3 | 0 | 0 |

**Table S4.** Unfolding times were estimated for two starting misfolded conformations (a Loss of native entanglement conformation; and a Gain of non-native entanglement conformation) at 298 K in all-atom simulations using super-Arrhenius analysis for gene product from gene ispE. Unfolding criteria is $Q \leq Q_{threshold}$ for 1 ns with various values of $Q_{threshold}$ to determine the unfolding time. Disentangling criteria is $G_0 + G_1 = 0$ for 1 ns. For details see SI Methods. Note well, the unfolding times listed have been corrected for the observed acceleration in unfolding rates in all-atom simulations.

| Misfolded state type | Criterion | Unfolding time 298 K (seconds) | 95% confidence interval (seconds) |
|---|---|---|---|
| Loss of native entanglement | $Q \leq Q_{threshold}$ for 1 ns | | |
| | 0.40 | $2.102\times10^{4}$ (~5.8 h) | [115.585, $4.610\times10^{6}$] |
| | 0.35 | $1.567\times10^{4}$ (~4.3 h) | [139.692, $1.827\times10^{6}$] |
| | 0.30 | $7.908\times10^{3}$ (~2.2 h) | [71.809, $1.092\times10^{6}$] |
| | 0.25 | $3.925\times10^{4}$ s (~10.9 h) | [359.220, $4.340\times10^{6}$] |
| | 0.20 | $6.289\times10^{4}$ s (~17.5 h) | [690.694, $6.281\times10^{6}$ ] |
| Gain of non-native entanglement | $G_0 + G_1 = 0$ for 1 ns | $2.413\times10^{6}$ s (~27.9 days) | [931.587, $4.338\times10^{9}$] |

**Table S5**: Radius of gyration and solubility of the native and entangled states of Ubiquitin, λ-repressor, and 4-Diphosphocytidyl-2-C-Methyl-D-Erythritol Kinase.

| ID | Radius of gyration ($R_g$, nm) | $R_g$ difference compared to native structure (%) | Solubility |
|---|---|---|---|
| Ubiquitin | | | |
| Native Structure | 1.18 | | |
| 1 | 1.24 | 5.3% | 0.836 |
| 2 | 1.19 | 0.9% | 0.915 |
| 3 | 1.21 | 2.6% | 0.937 |
| 4 | 1.30 | 10.6% | 0.840 |
| 5 | 1.23 | 4.3% | 0.897 |
| 6 | 1.25 | 6.6% | 0.928 |
| 7 | 1.23 | 4.8% | 0.926 |
| 8 | 1.23 | 5.1% | 0.849 |
| 9 | 1.25 | 6.2% | 0.946 |
| 10 | 1.24 | 5.5% | 0.908 |
| 11 | 1.21 | 3.1% | 0.903 |
| 12 | 1.20 | 1.7% | 0.940 |
| 13 | 1.20 | 2.2% | 0.990 |

| | | | |
|---|---|---|---|
| 14 | 1.20 | 2.0% | 0.955 |
| 15 | 1.22 | 3.6% | 0.903 |
| 16 | 1.30 | 10.8% | 0.901 |
| 17 | 1.26 | 7.3% | 0.963 |
| 18 | 1.26 | 6.8% | 0.787 |
| 19 | 1.26 | 7.1% | 0.958 |
| 20 | 1.31 | 11.2% | 0.976 |
| 21 | 1.27 | 7.7% | 0.807 |
| The average difference between entangled states and native structure | | 5.5% | |
| λ-repressor | | | |
| Native Structure | 1.19 | | |
| 22 | 1.25 | 5.7% | 0.802 |
| 23 | 1.25 | 5.4% | 0.832 |
| 24 | 1.21 | 2.2% | 0.770 |
| 25 | 1.25 | 5.3% | 0.739 |
| 26 | 1.27 | 7.5% | 0.747 |
| 27 | 1.27 | 7.3% | 0.755 |
| 28 | 1.24 | 4.9% | 0.729 |
| 29 | 1.25 | 5.8% | 0.673 |
| 30 | 1.21 | 2.0% | 0.766 |
| 31 | 1.26 | 6.1% | 0.710 |
| 32 | 1.26 | 6.1% | 0.719 |
| 33 | 1.24 | 4.4% | 0.765 |
| The average difference between entangled states and native structure | | 5.2% | |
| 4-Diphosphocytidyl-2-C-Methyl-D-Erythritol Kinase | | | |
| Native Structure | 1.93 | | |
| Loss of native entanglement | 2.08 | 8.2% | 0.942 |
| Gain of non-native entanglement | 1.97 | 2.4% | 1.0 |
| The average difference between entangled states and native structure | | 5.3% | |

The solubility of the entangled states was calculated using Eq. S8.

**Table S6**: Entangled state ID (table S2) and the frame corresponding to DE Shaw's trajectories.

| ID | Trajectory's names are from the D. E Shaw group as reported in Ref. [2,3] | Simulation Frame |
|---|---|---|
| Ubiquitin | | |

| | | |
|---|---|---|
| 1 | | 49551 |
| 2 | | 49552 |
| 3 | | 49553 |
| 4 | | 49601 |
| 5 | | 49548 |
| 6 | | 49517 |
| 7 | | 49519 |
| 8 | | 49544 |
| 9 | | 49599 |
| 10 | pnas2013-unfold-2-c-alpha-000.dcd | 49651 |
| 11 | | 49576 |
| 12 | | 49577 |
| 13 | | 49578 |
| 14 | | 49579 |
| 15 | | 49582 |
| 16 | | 49593 |
| 17 | | 49595 |
| 18 | | 49570 |
| 19 | | 49594 |
| 20 | pnas2013-native-4-c-alpha-006.dcd | 13525 |
| 21 | pnas2013-native-5-c-alpha-002.dcd | 48392 |
| λ-repressor | | |
| 22 | | 54713 |
| 23 | | 54729 |
| 24 | | 59891 |
| 25 | | 59863 |
| 26 | | 59878 |
| 27 | | 59698 |
| 28 | lambda-0-c-alpha-001.dcd | 59884 |
| 29 | | 59870 |
| 30 | | 59890 |
| 31 | | 59861 |
| 32 | | 59866 |
| 33 | | 59883 |

**Table S7:** Representative entanglements from each post-translational metastable state

| MSM state | native contact | crossing | $g_{N,ref}$ (Eq. S2) | $g_{N,state}$ (Eq. S2) |
|---|---|---|---|---|
| 0 | (33, 149) | 7 | -0.847 | 0.081 |
| 0 | (26, 185) | 10, 8 | -1.665 | 0.062 |
| 0 | (149, 276) | 33 | -0.085 | -0.7 |

| | | | | |
|---|---|---|---|---|
| 0 | (29, 196) | 16 | -1.032 | 0.62 |
| 1 | (32, 144) | 8 | -0.87 | 0.041 |
| 1 | (29, 196) | 17 | -1.032 | 0.727 |
| 1 | (26, 185) | 10, 8 | -1.665 | 0.092 |
| 1 | (150, 273) | 31 | -0.059 | -0.681 |
| 2 | (26, 185) | 10, 8 | -1.665 | 0.093 |
| 2 | (32, 144) | 8 | -0.87 | 0.065 |
| 2 | (29, 196) | 16 | -1.032 | 0.759 |
| 2 | (88, 122) | 46 | -0.264 | 0.637 |
| 3 | (88, 122) | 44 | -0.264 | 0.677 |
| 4 | (88, 122) | 44 | -0.264 | 0.611 |

**Table S8**: Overlaps of significant LiP-MS peptides with representative entanglements

| MSM state | Peptide cut site | Time point | Residues forming the native contact that closes the loop | Crossing residue | $g_{N,ref}$ (Eq. S2) | $g_{N,state}$ (Eq. S2) | Jaccard (Eq. S12) |
|---|---|---|---|---|---|---|---|
| 0 | G155 | 1h | (33, 149) | 7 | -0.847 | 0.081 | 0.152 |
| 0 | G155 | 1h | (149, 276) | 33 | -0.085 | -0.7 | 0.122 |
| **0** | **D210** | **10h** | **(29, 196)** | **16** | **-1.032** | **0.62** | **0.079** |
| **0** | **T161** | **10h** | **(149, 276)** | **33** | **-0.085** | **-0.7** | **0.022** |
| 0 | F32 | 10h | (33, 149) | 7 | -0.847 | 0.081 | 0.357 |
| 0 | F32 | 10h | (26, 185) | 10, 8 | -1.665 | 0.062 | 0.205 |
| 0 | F32 | 10h | (149, 276) | 33 | -0.085 | -0.7 | 0.122 |
| 0 | F32 | 10h | (29, 196) | 16 | -1.032 | 0.62 | 0.242 |
| **0** | **D210** | **24h** | **(29, 196)** | **16** | **-1.032** | **0.62** | **0.079** |
| **0** | **T161** | **24h** | **(149, 276)** | **33** | **-0.085** | **-0.7** | **0.022** |
| 1 | G155 | 1h | (150, 273) | 31 | -0.059 | -0.681 | 0.167 |
| 1 | D210 | 10h | (29, 196) | 17 | -1.032 | 0.727 | 0.026 |
| 1 | F32 | 10h | (32, 144) | 8 | -0.87 | 0.041 | 0.393 |
| 1 | F32 | 10h | (29, 196) | 17 | -1.032 | 0.727 | 0.258 |
| 1 | F32 | 10h | (26, 185) | 10, 8 | -1.665 | 0.092 | 0.196 |
| 1 | F32 | 10h | (150, 273) | 31 | -0.059 | -0.681 | 0.135 |
| 1 | D210 | 24h | (29, 196) | 17 | -1.032 | 0.727 | 0.026 |
| 2 | L118 | 1h | (88, 122) | 46 | -0.264 | 0.637 | 0.219 |
| 2 | D210 | 10h | (29, 196) | 16 | -1.032 | 0.759 | 0.079 |
| 2 | F32 | 10h | (26, 185) | 10, 8 | -1.665 | 0.093 | 0.2 |
| 2 | F32 | 10h | (32, 144) | 8 | -0.87 | 0.065 | 0.367 |
| 2 | F32 | 10h | (29, 196) | 16 | -1.032 | 0.759 | 0.242 |
| 2 | D210 | 24h | (29, 196) | 16 | -1.032 | 0.759 | 0.079 |

| | | | | | | | |
|---|---|---|---|---|---|---|---|
| 2 | [47-72] | 24h | (88, 122) | 46 | -0.264 | 0.637 | 0.039 |
| 3 | L118 | 1h | (88, 122) | 44 | -0.264 | 0.677 | 0.258 |
| 4 | L118 | 1h | (88, 122) | 44 | -0.264 | 0.611 | 0.216 |

#### References


1. Jorgensen, W. L., Chandrasekhar, J., Madura, J. D., Impey, R. W. & Klein, M. L. Comparison of simple potential functions for simulating liquid water. *J. Chem. Phys.* **79**, 926–935 (1983).

2. Piana, S., Lindorff-Larsen, K. & Shaw, D. E. Atomic-level description of ubiquitin folding. *Proc. Natl. Acad. Sci. U. S. A.* **110**, 5915–5920 (2013).

3. Lindorff-Larsen, K., Piana, S., Dror, R. O. & Shaw, D. E. How fast-folding proteins fold. *Science (80-. ).* **334**, 517–520 (2011).

4. Bryson, A. E. & Denham, W. F. A Steepest-Ascent Method for Solving Optimum Programming Problems. *J. Appl. Mech.* **29**, 247–257 (1962).

5. Haug, E. J., Arora, J. S. & Matsui, K. A steepest-descent method for optimization of mechanical systems. *J. Optim. Theory Appl.* **19**, 401–424 (1976).

6. Darden, T., York, D. & Pedersen, L. Particle mesh Ewald: An N·log(N) method for Ewald sums in large systems. *J. Chem. Phys.* **98**, 10089–10092 (1993).

7. Nosé, S. A unified formulation of the constant temperature molecular dynamics methods. *J. Chem. Phys.* **81**, 511–519 (1984).

8. William G. Hoover. Canonical dynamics: Equilibrium phase-space distributions William. *Phys. Rev. A* **31**, 1695–1697 (1985).

9. Parrinello, M. & Rahman, A. Polymorphic transitions in single crystals: A new molecular dynamics method. *J. Appl. Phys.* **52**, 7182–7190 (1981).

10. Hess, B., Bekker, H., Berendsen, H. J. C. & Fraaije, J. G. E. M. LINCS: A Linear Constraint Solver for molecular simulations. *J. Comput. Chem.* **18**, 1463–1472 (1997).

11. Abraham, M. J. *et al.* Gromacs: High performance molecular simulations through multi-level parallelism from laptops to supercomputers. *SoftwareX* **1–2**, 19–25 (2015).

12. Huang, J. *et al.* CHARMM36m: An improved force field for folded and intrinsically disordered proteins. *Nat. Methods* **14**, 71–73 (2016).

13. Jiang, Y. *et al.* How synonymous mutations alter enzyme structure and function over long time scales. *Nat. Chem.* (2022) In press.

14. Nissley, D. A. *et al.* Universal protein misfolding intermediates can bypass the proteostasis network and remain soluble and less functional. *Nat. Commun.* **13**, 3081 (2022).

15. Halder, R., Nissley, D. A., Sitarik, I. & O'Brien, E. P. Subpopulations of soluble, misfolded proteins commonly bypass chaperones: How it happens at the molecular level. *bioRxiv* (2021) doi:10.1101/2021.08.18.456736.

16. Niemyska, W., Millett, K. C. & Sulkowska, J. I. GLN: a method to reveal unique properties of lasso type topology in proteins. *Sci. Rep.* **10**, (2020).

17. Onuchic, J. N., Luthey-Schulten, Z. & Wolynes, P. G. THEORY OF PROTEIN FOLDING: The Energy Landscape Perspective. *Annu. Rev. Phys. Chem.* **48**, 545–600 (1997).

18. Sarkar, D., Kang, P., Nielsen, S. O. & Qin, Z. Non-Arrhenius Reaction-Diffusion Kinetics for Protein Inactivation over a Large Temperature Range #. *ACS Nano* **13**, 8669–8679 (2019).

19. Frishman, D. & Argos, P. Knowledge-based protein secondary structure assignment. *Proteins Struct. Funct. Bioinforma.* **23**, 566–579 (1995).

20. Tsolis, A. C., Papandreou, N. C., Iconomidou, V. A. & Hamodrakas, S. J. A Consensus Method for the Prediction of 'Aggregation-Prone' Peptides in Globular Proteins. *PLoS One* **8**, 1–6 (2013).

21. Gutierres, M. B. B., Bonorino, C. B. C. & Rigo, M. M. ChaperISM: Improved chaperone binding prediction using position-independent scoring matrices. *Bioinformatics* **36**, 735–741 (2020).

22. Moulder, R., Goo, Y. A. & Goodlett, D. R. Label-Free Quantitation for Clinical Proteomics. in *Quantitative Proteomics by Mass Spectrometry* (ed. Sechi, S.) 65–76 (Springer New York, 2016). doi:10.1007/978-1-4939-3524-6_4.

23. Veit, J. *et al.* LFQProfiler and RNPxl: Open-Source Tools for Label-Free Quantification and Protein-RNA Cross-Linking Integrated into Proteome Discoverer. *J. Proteome Res.* **15**, 3441–3448 (2016).

24. Bateman, A. *et al.* UniProt: the universal protein knowledgebase in 2021. *Nucleic Acids Res.* **49**, D480–D489 (2021).

25. To, P., Whitehead, B., Tarbox, H. E. & Fried, S. D. Nonrefoldability is Pervasive across the E. coli Proteome. *J. Am. Chem. Soc.* **143**, 11435–11448 (2021).

26. To, P., Xia, Y., Lee, S. O. Devlin, T., Fleming, K. G. & Fried, S. D. A proteome-wide map of chaperone-assisted protein refolding in a cytosol-like milieu. *Proc. Natl. Acad. Sci. USA*. (2022) In press .

27. Nissley, D. A. *et al.* Electrostatic Interactions Govern Extreme Nascent Protein Ejection Times from Ribosomes and Can Delay Ribosome Recycling. *J. Am. Chem. Soc.* **142**, 6103–6110 (2020).

28. Röblitz, S. & Weber, M. Fuzzy Spectral Clustering by PCCA+: Application to Markov State Models and Data Classification. *Adv. Data Anal. Classif.* **7**, 147–179 (2013).

29. Dabrowski-Tumanski, P., Rubach, P., Niemyska, W., Gren, B. A. & Sulkowska, J. I. Topoly: Python package to analyze topology of polymers. *Brief. Bioinform.* **22**, 1–8 (2021).
